# MusTer: A Generalizable Microfluidic Platform Combining Multi-Parametric Droplet Sorting and Multi-Droplet Merging in Single-Cell Sequencing

**DOI:** 10.64898/2025.12.29.696950

**Authors:** Luoquan Li, Weilun Liu, Guangyao Cheng, Fuyang Qu, Silin Zhong, Yi-Ping Ho

**Affiliations:** Department of Biomedical Engineering, The Chinese University of Hong Kong, Shatin, New Territories, Hong Kong SAR, China; State Key Laboratory of Agrobiotechnology, School of Life Sciences, The Chinese University of Hong Kong, Shatin, New Territories, Hong Kong SAR, China; State Key Laboratory of Marine Pollution, City University of Hong Kong, Hong Kong SAR, China; Hong Kong Branch of CAS Center for Excellence in Animal Evolution and Genetics, The Chinese University of Hong Kong, Hong Kong SAR, China; Centre for Novel Biomaterials, The Chinese University of Hong Kong, Shatin, New Territories, Hong Kong SAR, China

## Abstract

Droplet microfluidics is a core technology that powers high-throughput single cell sequencing. However, the current generation of single-cell microfluidics faces notable limitations, including cell aggregation, suboptimal on-chip reactions that compromise experimental outcomes and elevate background noise, as well as a dependence on costly commercial barcode beads. To address these challenges, we present MusTer, an integrated next-generation platform with multi-parametric singlet droplet sorting and triple-droplet merging capability. MusTer’s multi-parametric singlet sorting module enables in-line droplet analysis of intrinsic fluorescence peak amplitude, width and interval from single-nucleus- (singlet) or multiple-nuclei (multiplet)-encapsulating droplets, subsequently allowing an effective separation of the singlet droplets from multiplet droplets and empty droplets. MusTer’s triple-droplet merging module enables precise multi-step reactions, with each step performed under its own optimal conditions, thereby significantly enhancing experimental flexibility and efficiency. We validated MusTer’s performance by performing single-cell ATAC-seq on maize leaves. The results demonstrate that MusTer significantly reduces the doublet rate, enhances the signal-to-noise ratio, and yields improved cell clustering compared with traditional methods. These results validate MusTer’s capability to overcome key limitations in droplet-based single-cell analysis, effectively enhancing data quality and reliability, and also paves the way for its use in other challenging sample types and multi-step single-cell assays.

## Introduction

Single-cell omics has shown great promise in uncovering biological mechanisms, aiding in drug discovery, disease treatment and prognosis determination^1–6^. For example, single-cell transcriptome sequencing (scRNA-seq) maps the transcriptional profiles of individual cells at high resolution, facilitating the discovery of unique cell types, identification of abnormally expressed genes, illustration of developmental trajectories and dissection of the tumor microenvironment (TME)^7–10^. Single-cell assay for transposase-accessible chromatin using sequencing (scATAC-seq) enables the examination of chromatin accessibility across individual cells. It can uncover how variation in open chromatin regions regulates gene expression, facilitating the identification of cell subtypes, rare single nucleotide polymorphisms (SNPs), and copy number variations (CNVs)^11–13^. Conventional methods for single-cell analysis, such as isolating individual cells into tubes or sorting them into PCR plates, are labor-intensive and severely limited in throughput^14–17^. Whereas, droplet microfluidics has revolutionized the field by automating this process, enabling the high-throughput processing of thousands of single cells simultaneously while drastically reducing reagent consumption and manual labor^18–20^. Despite these advantages, achieving the precise encapsulation of a single cell or nucleus within an individual droplet (forming a singlet droplet) remains a critical technical challenge. The co-encapsulation of multiple cells or nuclei, resulting in doublet or multiplet droplets, makes the subsequent bioinformatic disambiguation difficult or impossible, often leading to irretrievable data loss^21, 22^.

Commonly seen droplet microfluidics-based platforms like the 10X Chromium rely on Poisson-limited dilution to minimize doublet formation^23^. This method is inherently inefficient, achieving a doublet rate of ∼8% in the final sequencing library comes at the cost of ∼90% wasted empty droplets during library preparation^24^. This doublet issue is even aggravated for the case of scATAC-seq, as the tagged nuclei are prone to aggregation^25, 26^, reducing the fraction of singlet droplets and thus the usable sequencing data. Experimental measures such as sample-based cell hashing^27, 28^ or combinatorial indexing can, to a certain extent, tolerate cell or nucleus co-encapsulation by introducing multiple rounds of barcoding^25, 29, 30^. However, sample-based cell hashing is limited to identifying cross-sample multiplets by assigning each cell into its sample of origins; cell clusters or aggregates that share the same sample index and are encapsulated within a single droplet remain indistinguishable. Similarly, although combinatorial indexing increases barcode diversity, it does not resolve pre-existing physical aggregates. For example, in scATAC-seq, nuclei that aggregate after tagmentation are labeled as a single unit and consequently carry identical barcode combinations, rendering the individual nuclei within the aggregate undifferentiable. Other methods, such as fluorescence-activated cell sorting (FACS) may physically remove cell/nucleus aggregates prior to droplet encapsulation^31, 32^. However, FACS introduces mechanical stress at prolonged duration, which may compromise cells/nuclei integrity and lead to DNA leakage or contamination^33^, which is a concern in scATAC-seq when performing FACS after tagmentation^31, 34^. Conversely, when FACS is performed before tagmentation, nuclei that are initially sorted as singlets may re-aggregate during the subsequent transposition process^25, 35^. As a result, aggregation-induced co-encapsulation of multiple nuclei into a single droplet remains unavoidable in scATAC-seq despite the use of FACS. Fluorescence-activated droplet sorting (FADS), such as in the workflow of spinDrop, can enrich the singlet droplets by excluding the empty droplets^36^. However, sorting solely based on fluorescence intensity may not effectively differentiate singlet from doublet or multiplet droplets. On the other hand, computational strategies, such as Scrublet^37^, scDbIFinder^38^, AMULET^39^, SnapATAC^40^ and ArchR^41^ are developed to identify and remove multiplets post-sequencing by comparing data across different cells to simulate multiplets generated from the available data and quantifying the number of aligned reads overlapping genomic regions. These simulation-based approaches, however, are shown ineffective, particularly exhibiting low precision in identifying multiplets originating from the same cell type^39^ and polyploid cells commonly found across most plant tissues^42^.

While droplet-based single-cell analysis substantially improves throughput compared to plate-based techniques, the droplet format inherently constrains the implementation of multi-step reactions, as reagent exchange within droplets is difficult to achieve. Consequently, sequential biochemical steps are often performed under suboptimal, compromised conditions within a single reaction environment. This limitation poses a major challenge for the development and optimization of complex multi-step protocols that are essential for many advanced single-cell omics applications^43–45^. For example, in scRNA-seq, cell lysis and reverse transcription are confined to the same droplet environment. The inability to optimize these steps independently is a key factor contributing to limited RNA capture efficiency^19, 46^. Similarly, in scATAC-seq, failure to inactivate Tn5 transposase before droplet encapsulation into PCR buffer leads to inefficient PCR amplification^47–49^. Techniques such as pico-injection can partially address these constraints by enabling reagent addition into flowing droplets^36^. However, pico-injection is restricted to the delivery of homogeneous solutions and cannot introduce insoluble components like cells or barcoded beads. Despite rapid advances in buffer exchange using microcapsule-based single-cell omics technologies, persistent challenges related to contamination remain^50,51^. Addressing these concerns necessitates precise tuning of the microcapsule pore size to ensure compatibility with the sequencing of specific target products.

To overcome these limitations of existing droplet-based single-cell sequencing platforms, we developed **MusTer**, a microfluidic system that integrates **mu**lti-parametric **s**inglet droplet sorting with **t**riple-droplet m**er**ging. MusTer incorporates an on-chip multi-parametric sorting module designed to selectively remove multiplet droplets and empty droplets, thereby enriching true singlet droplets prior to downstream processing. Compared to above-mentioned experimental strategies on eliminating multiplets, the sorting module in MusTer is particularly advantageous for samples prone to aggregates, while preserving the integrity of singlets. Following sorting, a triple-droplet merging module combines individual sorted singlet droplets with droplets containing barcode templates and droplets containing PCR reagents, enabling multi-step reactions to be performed at optimal conditions inside droplets.

We demonstrate the effectiveness of MusTer by applying it to scATAC-seq of maize (*Zea mays*) as a model plant species. Maize leaves employ a unique C4 photosynthetic pathway, which enhances water-use efficiency even under high-temperature and drought conditions^52, 53^. This complex pathway requires tight cooperation of transcriptional regulation between two distinct cell types: mesophyll and bundle sheath cells, making it well suited for evaluating chromatin accessibility at single-cell resolution^54^. However, scATAC-seq for maize leaves remains a challenging task due to nuclei aggregation during extraction and Tn5 tagmentation. Previous scATAC-seq methods have been constrained by a trade-off between throughput and data quality, as exemplified by the high throughput droplet-based 10X Chromium system^55^ versus the superior data resolution of the low-throughput plate-based mthods^30,56^. Here in this study, by profiling chromatin accessibility in 9,120 maize nuclei, we show that MusTer’s on-chip singlet droplets sorting module dramatically reduced doublet rates, while its triple-droplet merging module significantly improved data quality compared to traditional microfluidic systems^55^.

## Results

### Workflow of MusTer scATAC-seq

Targeting maximized utilization of sequencing data and enhanced reaction efficiency in droplet-based single-cell omics, MusTer was designed to use on-chip multi-parametric sorting to identify and enrich singlet droplets, which were then merged with a pair of reagent droplets for barcoding and amplification. To demonstrate its capabilities, we applied MusTer to droplet-based scATAC-seq of maize (**Fig. 1**), a challenging application due to nuclei aggregation during extraction and Tn5 tagmentation, as well as suboptimal reaction efficiencies in conventional one-pot workflows. In the presented workflow, the maize B73 leaf nuclei were extracted, tagmented by Tn5 transposase to label the open chromatin regions, and stained with SYBR Green I. Thermolabile proteinase K were added to the nuclei during droplets formation, and the nuclei droplets were subsequently introduced to the multi-parametric singlet droplet sorting module (**Fig. 1**). Droplets containing a single nucleus is selected based on a series of criteria, such as the peak amplitude, peak width and peak interval. Thermolabile proteinase K in the sorted singlet droplets was then activated to denature the Tn5 proteins by incubating the droplets at 37°C prior to the barcoding and amplification step.

**Fig. 1.**
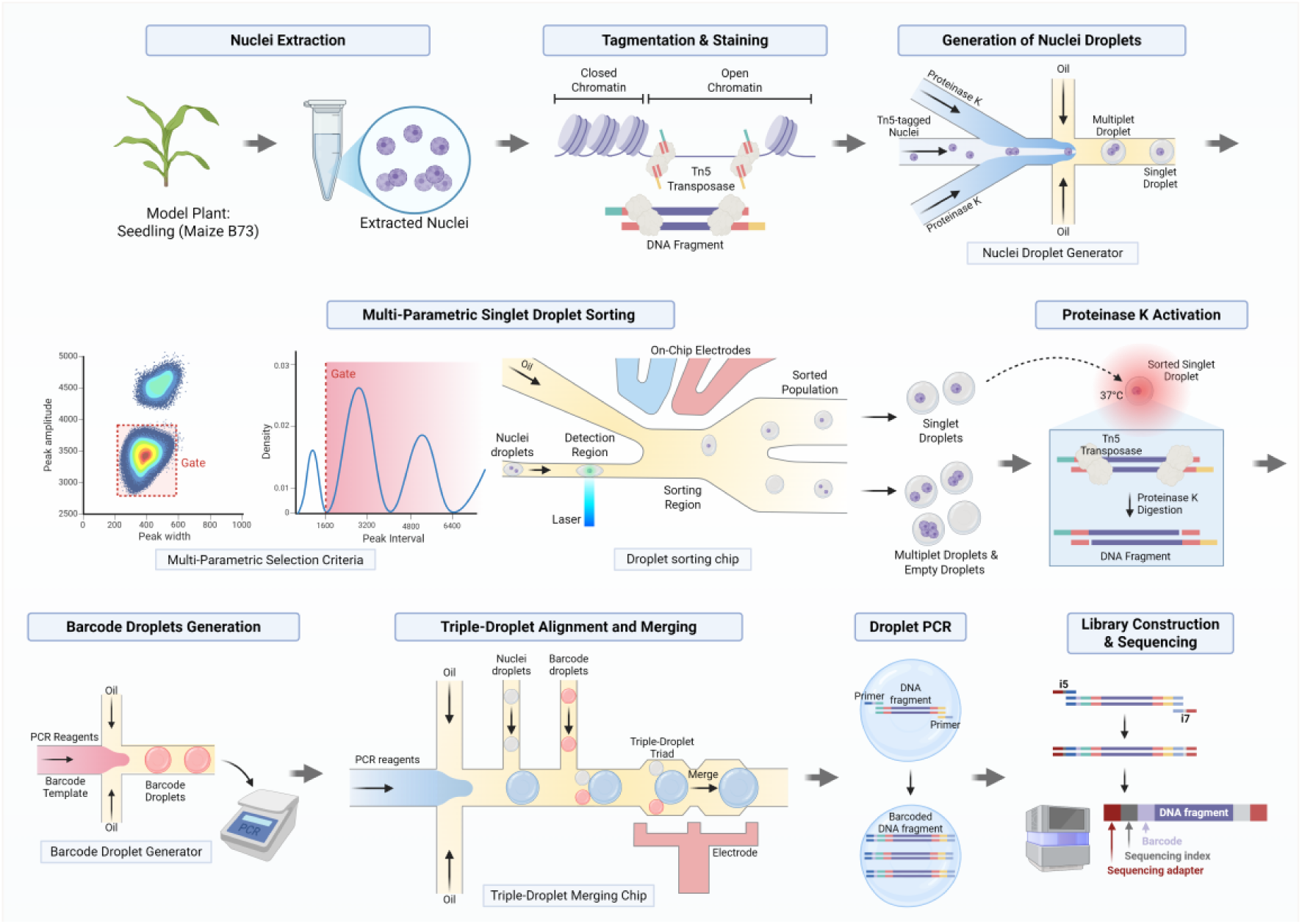
Workflow of MusTer scATAC-seq. Selected as a model, nuclei were extracted from Maize (*Zea mays* B73 as a model) seedlings by the chopped leaves. Nuclei open chromatin DNA was fragmented by Tn5 transposase tagmentation and stained by SYBR Green I. Nuclei droplets were generated by co-encapsulating tagged nuclei and thermolabile proteinase K using the nuclei droplet generator (**Fig. S1a**). Multi-parametric singlet droplet sorting was performed on the droplet sorting chip (**Fig. S2**) embedded with a set of electrodes to identify and separate the singlet droplets from those empty and multiplet droplets. After sorting, proteinase K within the singlet droplets was activated at 37°C to denature Tn5 proteins on the open chromatin fragments. A collection of barcode droplets was separately generated through droplet digital PCR using the barcode droplet generator (**Fig. S1b**). The triplet-droplet merging module was designed to produce the PCR droplets encapsulating PCR reagents, and to match the generated PCR droplet with a reinjected sorted singlet droplet and a barcode droplet, forming a triple-droplet triad with the triple-droplet merging chip (**Fig. S3**). The triple-droplet triad was merged by dielectrophoretic force on-chip. Droplet PCR was performed to amplify the tagged open chromatin DNA with specific barcode primer. Upon breaking the droplets, another round of bulk PCR was performed to introduce sequencing index i5, i7 and sequencing adapters. The produced library was purified for next generation sequencing. Fig.1 is created with BioRender.com.

Instead of relying on commercial or custom-synthesized barcode beads, such as hydrogel^23^ ^47, 57^, polystyrene or silica beads^18, 58, 59^, we employed a validated droplet-based single-cell indexing method^20^ that can be readily implemented in labs equipped with standard microfluidics. In this approach, droplets containing specific barcode templates (hereafter termed the barcode droplet, **Fig. 1**) are generated and amplified by droplet digital PCR (ddPCR)^60^, providing a simple and scalable means of barcode generation. Although droplet-based indexing may exhibit a lower effective barcoding rate compared to bead-based approaches, it substantially reduces experimental complexity and eliminates the need for specialized bead synthesis. Importantly, because MusTer incorporates a droplet merging module for barcode delivery, the platform is inherently compatible with alternative barcoding schemes such as barcode beads, if preferred.

Following Tn5 inactivation, the triple-droplet merging module was used to fuse a barcode droplet with a sorted singlet droplet, and a droplet containing PCR reagents (hereafter termed the PCR droplet, **Fig. 1**), followed by an *in situ* droplet PCR to generate barcoded ATAC-seq libraries. The droplets were then broken, and DNA was purified for another round of PCR to add sequencing adapters for Illumina NGS sequencing.

### Characteristics of Droplets Containing Different Numbers of Nuclei

To characterize droplets containing nuclei, we employed a customized FADS setup in the multi-parametric singlet droplet sorting module (detailed in **Methods** and **Fig. S4**) for in-line droplet analysis and sorting. Considering the requirement for uniform laser excitation of nuclei within droplets and the default routing of untriggered droplets to the non-target outlet, the droplet detection and sorting regions were designed as shown in **Fig. 2a**. To fully cover the entire detection region, the laser beam was optimally elongated to around 100 µm, 4 times of the detection region channel width (25 µm) (**Fig. S5**). The focal point of the beam was positioned at the center of the channel widthwise to ensure a uniform excitation on the flowing droplets. A custom FPGA module was implemented on the Zynq programmable logic (PL) core to process the raw fluorescence data on-the-fly. Droplets were characterized based on the amplitude, width and interval of acquired fluorescence peaks as illustrated in **Fig. 2b**: The peak amplitude extracted the maximum recorded fluorescence intensity, mainly reflecting the fluorophore concentration and size of the fluorescent objects; The peak width registered the temporal duration, revealing the size and flow velocity of the fluorescent objects; The peak interval noted the time between consecutive peak termination events, collectively determined by the distance and flow velocity of fluorescent objects. All parameters were updated for in 1 µs post-peak termination (clock rate: 1 MHz) to ensure short decision latency.

**Fig. 2.**
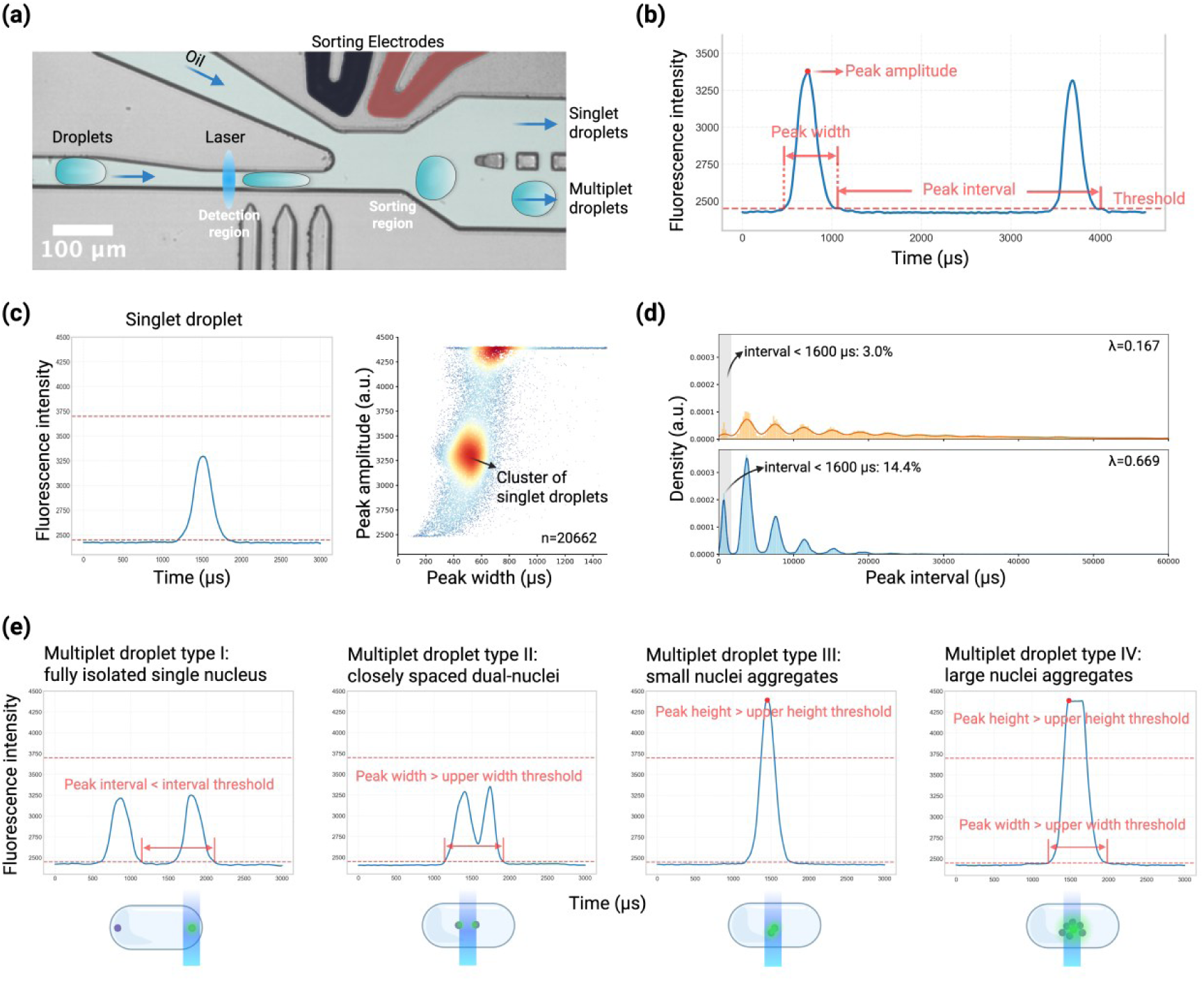
Signal Characteristics of Droplets Containing Nuclei Singlet and Multiplet. (a) Pseudo-colored microscopic image of the droplet detection and sorting region in the droplet sorting chip. The laser beam was optimally configured and expanded to 100 µm widthwise, allowing a uniform illumination of the entire channel width (25 µm). Fluorescence signals were recorded as W/O droplets passing through the beam, following by a triggered sorting at the sorting region based on the signal characteristics. (b) Representative fluorescence time trace from the detection. A threshold (dashed line) was used to identify valid peaks in real-time using a FPGA module. Key features, including the peak amplitude (maximum signal intensity), peak width (duration above the threshold), and peak interval (time between the end of two consecutive peaks), were extracted. (c) Signal characteristics of a singlet droplet. Left: A single fluorescent peak with amplitude confined between 3,000-3,700 a.u., and a width between 200-600 µs, separated from other peaks of more than 1,600 at the operated flow rate. Right: Amplitude-width scatter plot was constructed from 20,662 analyzed peaks, where the events of singlet droplets were clustered densely in the center. (d) Peak interval distributions of two droplet populations with different proportions of Type I multiplet droplets indicated by the Poisson expectation parameter (λ). Both distributions were observed periodic corresponding to the inter-droplet spacing. The population of higher λ (lower panel, λ = 0.669) presented an elevated fraction of sub-1,600 µs intervals (14.4%), indicating more peaks from isolated nuclei within the same droplet (Type I multiplet droplet). (e) Representative fluorescence traces and illustrations of the four major multiplet droplets. **Type I, Fully Isolated Single Nucleus**: Multiple narrow and non-overlapping peaks with intervals below 1,600 µs. **Type II, Closely Spaced Dual-Nuclei**: Single broadening peak exceeding the upper width threshold. **Type III, Small Aggregates:** Peaks with amplitudes above the upper height threshold and similar peak width as the singlet droplets. **Type IV, Large Aggregates**: Broad and high-amplitude peaks exceeding the thresholds of both width and height. A multi-parametric criteria was developed accordingly to distinguish single droplets from the four major types of multiplet droplets and multiplet droplets of combined features.

Droplets containing different numbers of extracted maize nuclei were examined and five different characteristic peaks were observed as the following. At a flow rate of 7 µL/min (equivalent to ∼14,000 droplets/min), singlet droplets were characterized with a narrow peak (width: 200-600 µs) separated from each other by at least 1,600 µs, which is the duration required for a droplet to pass through the laser beam. By modulating the gain voltage of the photomultiplier tube (PMT), the peak amplitude of singlet droplet was measured from 3,000 to 3,700 (a.u.: arbitrary unit), allowing an illustration of singlet droplets as a concentrated cluster on the amplitude-width scatter plot (**Fig. 2c**). Additionally, four main distinct multiplet droplets (Types I-IV) were identified based on the spatial distribution of nuclei observed from the bright field image and the corresponding temporal transit profiles through the laser beam (**Fig. 2e**, **Supplementary Movie 1**). Type I multiplet droplets, arose from fully isolated nuclei, generated sequential and non-overlapping peaks with peak intervals shorter than the droplet transit time. The peak *per se* of Type I multiplet droplets were observed similar to those from singlet droplets, therefore a measure of the peak interval was necessary to distinguish the two. The distribution of peak interval was plotted for two droplet populations with different proportion of Type I multiplet droplets in **Fig. 2d**. The peak interval showed a period distribution, where the period matched with the gap of droplets passing through the laser beam. The percentage of peak interval below 1,600 µs were observed proportional to the percentage of Type I multiplet droplets, proving the capability to identify Type I multiplet droplets by the peak intervals. Therefore, setting a minimum threshold of the peak interval for sorting was potentially capable of excluding Type I multiplet droplets. Type II multiplet droplets, characterized by closely spaced nuclei, exhibited extended peak widths due to overlapping laser transits (**Fig. 2e**), which could be filtered *via* an upper threshold on the peak width. On the other hand, multiplet droplets containing different level of nuclei aggregates generated signals as shown in Type III and Type IV, where the fluorescent peak height was observed stronger than singlet droplets and different peak width based on the level of aggregation. Type III multiplet droplets from sub-beam-size nuclei aggregates producing amplitude-enhanced peaks of a width similar to that of singlet droplets (**Fig. 2e**) were excluded through an upper threshold on the peak amplitude. Type IV multiplet droplets, formed by supra-beam-size nuclei aggregates, were observed with peaks of both increased width and amplitude (**Fig. 2e**), and were removed using either upper threshold for peak width or amplitude. A multi-parametric criteria was then established according to the above-mentioned signal characteristics to distinguish the singlet droplets from multiplet droplets. Notably, the employed criteria would also presumably exclude multiplet droplets with combined features of the four major types of multiplet droplets effectively.

### Validation of the Multi-Parametric Sorting Module by Isolating Fluorescent Mimics

To validate the proposed multi-parametric criteria for the isolation of singlet droplets, we conducted a series of experiments using fluorescein isothiocyanate (FITC)-labeled polystyrene (PS) microbeads. Given the size similarity^61^, 4-µm and 10-µm PS beads were employed to mimic the size of single plant nucleus and large nuclei aggregates, respectively. Firstly, droplets containing either 4-µm (mimics of singlet droplets) or 10-µm beads (mimics of Type IV multiplet droplets) were separately prepared. The two droplet populations were then mixed and injected into the droplet sorting chip. The PMT gain voltage was adjusted to allow for the amplitude of fluorescence peaks corresponding to singlet droplets distributing around 3,000-3,700 a.u. and clustered in the center on the width-amplitude plot, whereas the peak amplitude for 10-µm beads appeared saturated (**Fig. 3a Left**). Under this setting, a gating around the singlet droplet cluster (dashed boxed) was applied to allow for droplets meeting the selection criteria being sorted by the activated DEP force at the sorting region. Successful selection of droplet containing a 4-µm bead and exclusion of droplet containing a 10-µm bead was visualized in the image sequences in **Fig. 3b** extracted from **Supplementary Movie 2**. Representative images (**Fig. 3a Middle**) and quantifications (**Fig. 3a Right**) showed an enrichment of droplets containing single 4-µm bead (62.8% v.s. 99.7% before and after the sorting), validating that the combined selection criteria using the peak amplitude and width may effectively exclude Type IV multiplet droplets. It is important to note that the reported percentages were normalized to droplets containing detectable fluorescent beads excluding the empty droplets.

**Fig. 3.**
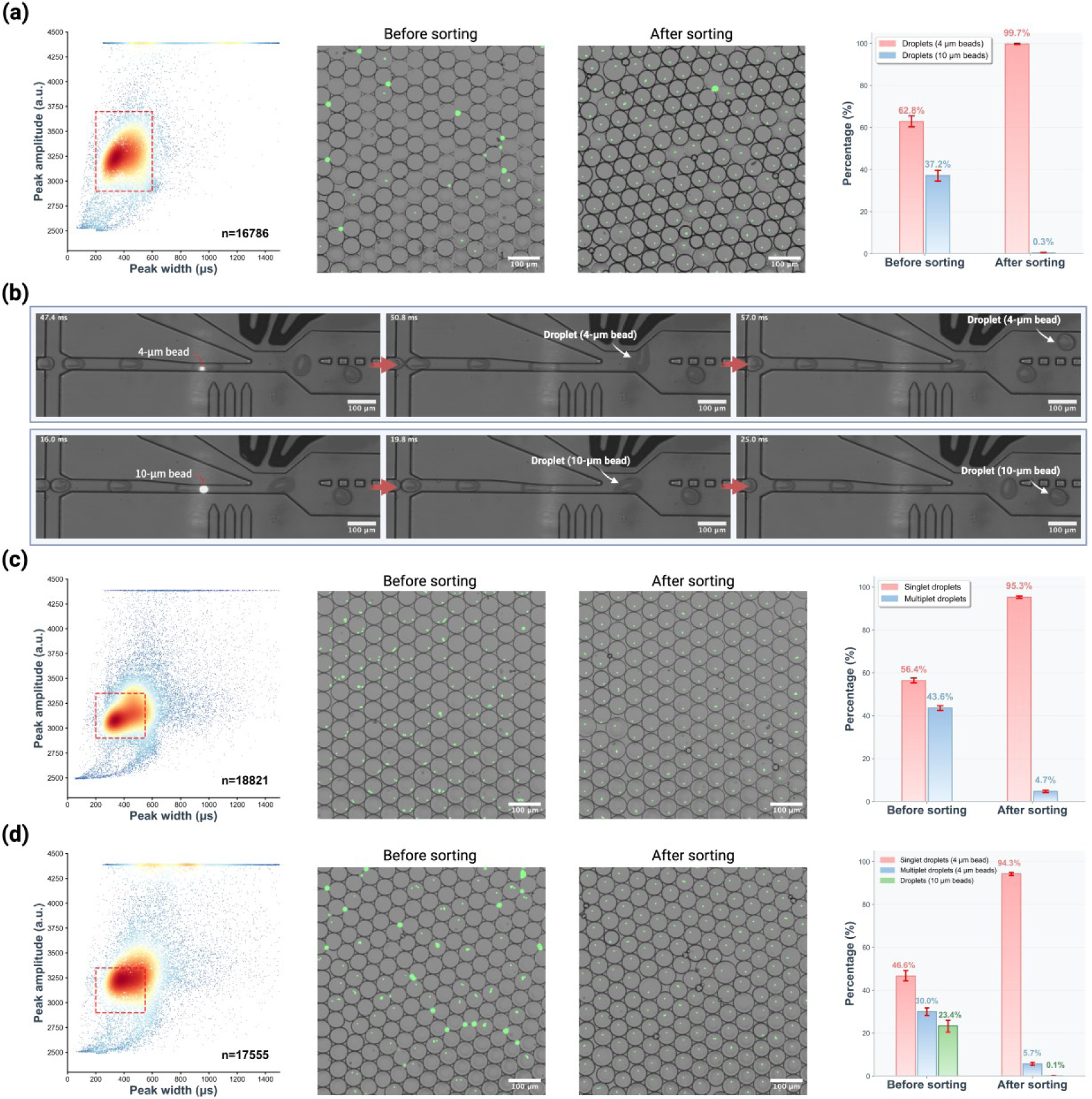
Validation of the Multi-Parametric Selection Criteria in Enriching Singlet Droplets Using Fluorescent Mimics. (a) Exclusion of Type IV multiplet droplets. Droplets containing either 4-µm or 10-µm fluorescent PS beads were used to simulate singlet droplets and Type IV multiplet droplets, respectively. Left: Amplitude-width scatter plot showing distinct droplet populations, with the singlet droplet cluster gated in the red dashed box. Due to limited PMT dynamic range, peaks from droplet containing 10-µm beads appeared saturated. Middle: Microscopic images of droplets before and after sorting, demonstrating effective removal of droplets containing 10-µm beads. Right: Quantification indicated that the fraction of droplets containing 4-µm beads increased from 62.8% to 99.7% after sorting, while droplets containing 10-µm beads reduced from 37.2% to 0.3%. (b) Representative image sequences (extracted from **Supplementary Movie 2**) showing the process of sorting droplets containing 4-µm bead and rejecting droplets containing 10-µm bead. (c) Exclusion of Type I-III multiplet droplets. A high-concentration suspension of 4-µm beads was used to generate droplets mimicking both singlet droplets and Type I-III multiplet droplets. Left: Scatter plot showing the gated singlet population (red dashed box). A fixed minimum peak interval threshold of 1,600 µs was applied. Middle: Microscopic images of droplets before and after sorting show effective removal of multiplet droplets. Right: The proportion of singlets droplets increased from 56.4% to 95.3% after sorting, while multiplet droplets decreased from 43.6% to 4.7%. (d) Validation using a complex droplet mixture containing singlet droplets and all multiplet droplets. Left: Scatter plot shows the composite population, with the gating condition indicated by the red dashed box. Minimum peak interval threshold was fixed at 1,600 µs. Middle: Microscopic images of droplets before and after sorting revealed effective enrichment of the singlet droplets. Right: The fraction of singlet droplets increased from 46.6% to 94.3%, while Type I-III and Type IV multiplet droplets were reduced to 5.7% and 0.1%, respectively. All percentages were normalized to bead-containing droplets, excluding empty droplets. Error bars expressed as 95% confidence interval of bootstrap resampled distribution from the acquired images, where droplet number N>2,000 for each data after sorting.

The selection criteria were also evaluated against Type I-III multiplet droplets by adjusting the concentration of 4-µm beads during droplet generation to modulate the average number of beads *per* droplet, mimicking the situations of multiple nuclei presented in one droplet with different proximity (Type I and II) and small aggregates (Type III). As shown in **Fig. 3c** and **Supplementary Movie 3**, the multi-parametric selection criteria, constraints on peak amplitude, width, and interval (minimum threshold fixed at 1,600 µs), were shown effective to enrich the population of droplets containing single 4-µm bead from 56.4% to 95.3%. Taken together, the multi-parametric selection criteria is expected effective in separating the singlet droplets from the droplets containing spatially separated beads (Type I), partially overlapping beads (Type II), and small bead aggregates (Type III), while the stringency of gating is tunable in balancing the isolation purity and recovery rate (**Fig. S6**).

The selection was then validated against complex conditions, wherein all four multiplet droplet mimics were in presence. Droplets containing 10-µm beads (Type IV multiplet droplet mimics) and various numbers of 4-µm beads (singlet droplet and Types I-III multiplet droplet mimics) were prepared. From **Fig. 3d**, 46.6% of droplets containing single 4-µm bead were identified before sorting. The fraction increased to 94.3% after sorting by applying the multi-parametric selection criteria, with all classes of multiplet droplet mimics substantially depleted (**Fig. 3d**, **Supplementary Movie 4**).

These results have prepared sufficient confidence for the application of proposed multi-parametric selection criteria to exclude multiplet droplets from the singlet counterparts.

### Implementation of the Multi-Parametric Sorting for Nuclei Droplets in scATAC-seq

Based on validation with fluorescent mimics, the multi-parametric selection criteria were then applied to sort singlet droplets with real nuclei samples post-transposition. Nuclei were extracted from maize B73 leaves, tagged with Tn5 transposase, and stained with SYBR Green I. Droplets were then produced at a nuclei concentration of ∼3,500/μL (Poisson parameter λ ∼ 0.24). Tn5 tagmentation was observed to aggravate the aggregation issue of the extracted nuclei (**Fig. S7**) as previously reported^25, 26^, generating a population with around 41.2% singlet droplets, 1.2% Type I-II multiplet droplets, and 57.6% Type III-IV multiplet droplets before sorting (**Fig. 4a** and **4d**). The selection of nuclei singlet droplets was further optimized by tuning the gating. Briefly, the presence of singlet droplets population on the amplitude-width scatter plot was regulated to allow sufficient distinction from other populations. Given that the fluorescence intensity serves as the major difference between the singlet droplets population and others, the PMT gain voltage was adjusted within the range of 0.56-0.6V. As shown in **Fig. 4b** and **4c**, increasing the PMT gain, i.e. the overall peak amplitude, was observed to curtain off the singlet droplet population from others by moving the events further apart from each other. At the PMT gain of 0.6V, gating as indicated by the red dashed box retained 32.3% of singlet droplet events among all events (corresponding to ∼78.4% recovery rate). Together with a fixed minimal peak-interval threshold at 1,600 µs, the gating (**Supplementary Movie 5**) presented an efficient enrichment of singlet droplets up to ∼95.0% (**Fig. 4b Right, Fig. 4d**). As a comparison, at 0.56 V of PMT gain voltage, the population of singlet droplet was not sufficiently differentiable with the populations of droplets containing presumably nuclei doublets or small aggregates with slightly higher peak amplitude than the singlet (**Fig. 4c Left**). Gating, in this case, accounted 51.1% of events from all, showed an insufficient separation of singlet droplets from multiplet droplets (75.7% and 24.3% of singlet and multiplet droplets, respectively as quantified in **Fig. 4c Right** and **Fig. 4d**). Consistent with the fraction of singlet droplets observed before sorting (41.2%), more than 50% of gated events would naturally include part of the multiplet droplets. Furthermore, these results also suggest that selection by fluorescence intensity alone is unable to fully isolate the singlet droplets, particularly against those containing doublets or small aggregates.

**Fig. 4.**
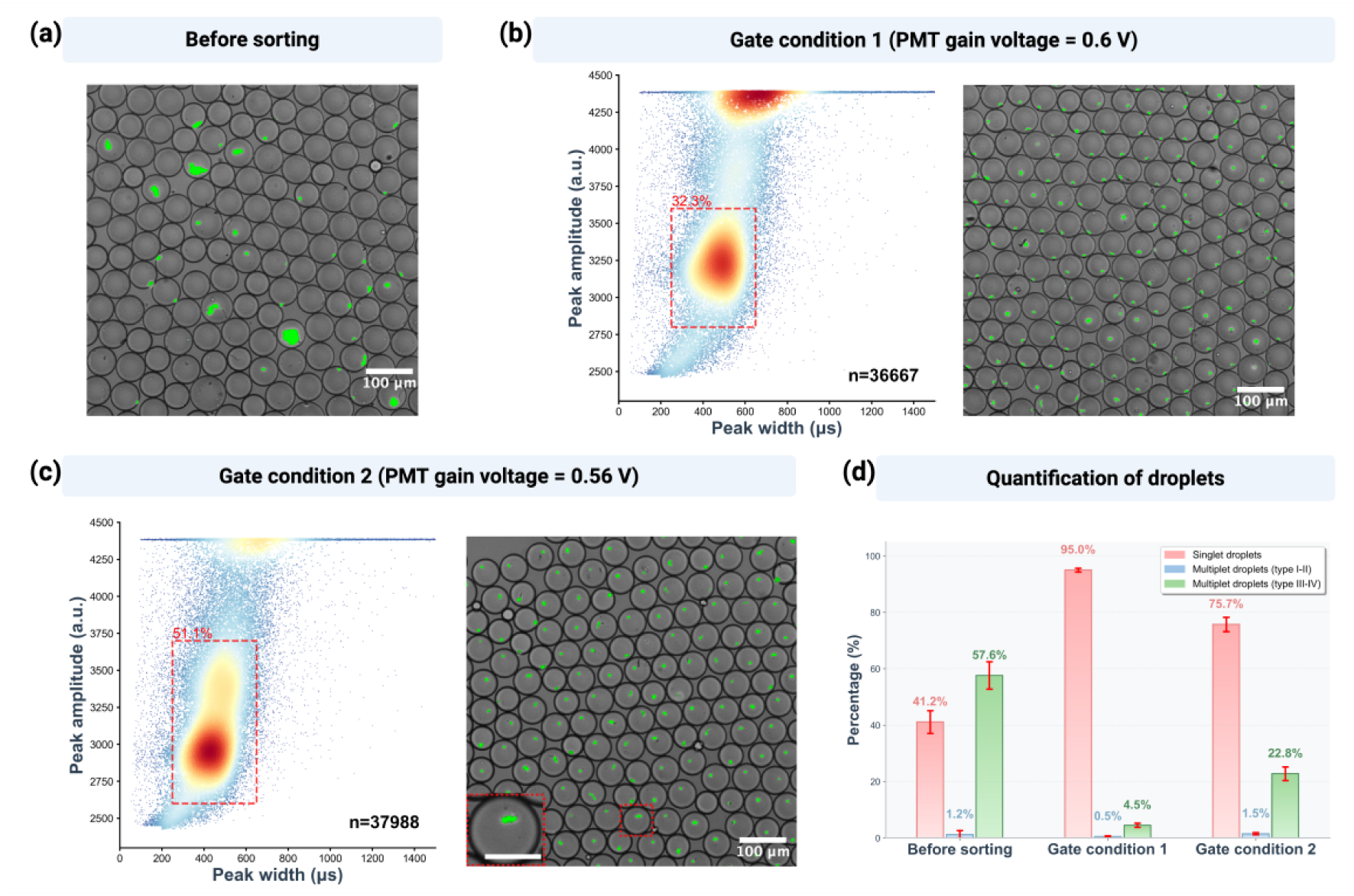
Enrichment of Singlet Droplets Encapsulating Nucleus with the Multi-Parametric Selection Criteria. (a) Microscopic image of nuclei droplets encapsulating tagged nuclei before sorting. Scale bar: 50 µm. (b) Left: Amplitude-width scatter plot at 0.6 V of PMT gain voltage with the gating condition 1 indicated by the red dashed box for sorting. Right: Microscopic image of nuclei droplets after sorting with gating condition 1. Scale bar: 50 µm. (c) Left: Amplitude-width scatter plot at 0.56 V of PMT gain voltage with the gating condition 2 indicated by the red dashed box for sorting. Right: Microscopic image of nuclei droplets after sorting with gating condition 2. The inset at bottom left showed a magnified view of nuclei multiplets encapsulated in a droplet. Scale bar: 50 µm. (d) Quantification indicated that the fraction of singlet droplets was enriched from 41.2% to 95% with gate condition 1 and 75.7% with gate condition 2. Minimum peak interval threshold was fixed at 1,600 µs. The percentage of multiplet droplets (type III-IV) was quantified as detailed in **Fig. S8**. Error bars expressed as 95% confidence interval of bootstrap resampled distribution from the acquired images, where droplet number N>3,000 for each data after sorting.

In addition, the proportion of empty droplets after the sorting was observed below 3%, minimizing the influence of background noise of final sequencing data originating from the empty droplets. The capability and reliability of the multi-parametric selection criteria to enrich singlet droplets was further validated at a higher nuclei concentrations during droplet generation (λ ∼ 0.95), where higher percentage of Type I multiplet droplets (separated nuclei) were presented. After sorting, the percentage of singlet droplets was increased from 41.9% to 96.9% (**Fig. S9**), showing the reliability of the proposed multi-parametric selection criteria under various conditions, and the potential of higher processing throughput with much less empty droplets (38.7% at λ ∼ 0.95 compared with 78.7% at λ ∼ 0.24).

### Triple-Droplet Merging Module in MusTer for scATAC-seq

In conventional droplet-based single-cell assays, suboptimal reaction efficiency is often tolerated, leading to compromised data quality. Droplet merging presents a promising strategy to overcome this limitation by enabling separate multi-step reactions within individual droplets^36^. Therefore, we designed a multi-droplet merging module for MusTer. In scATAC-seq, amplification efficiency was shown altered by the binding of Tn5 proteins after transposition^62–64^. Sodium dodecyl sulfate (SDS, 0.05% v/v) and proteinase K (1%, v/v), previously reported effective treatments^34, 65, 66^, were therefore introduced to deactivate Tn5 proteins post-tagmentation. Real-time quantitative PCR (qPCR) results shown in **Fig. 5a**, demonstrated comparable amplification efficiencies (Cycle Threshold (Ct) value of ∼12.78 and ∼12.88 for the SDS- and proteinase K-treated groups, respectively), and yet improved efficiency compared to the untreated group (Ct ∼ 15.56). However, droplets containing SDS treated samples were observed unstable (**Fig. S10)**. Consequently, proteinase K was selected for Tn5 proteins inactivation for the subsequent validation by co-encapsulation with the Tn5-tagged, SYBR-stained nuclei (λ ∼ 0.24). The proteinase K was activated following nuclei droplets sorting.

**Fig. 5.**
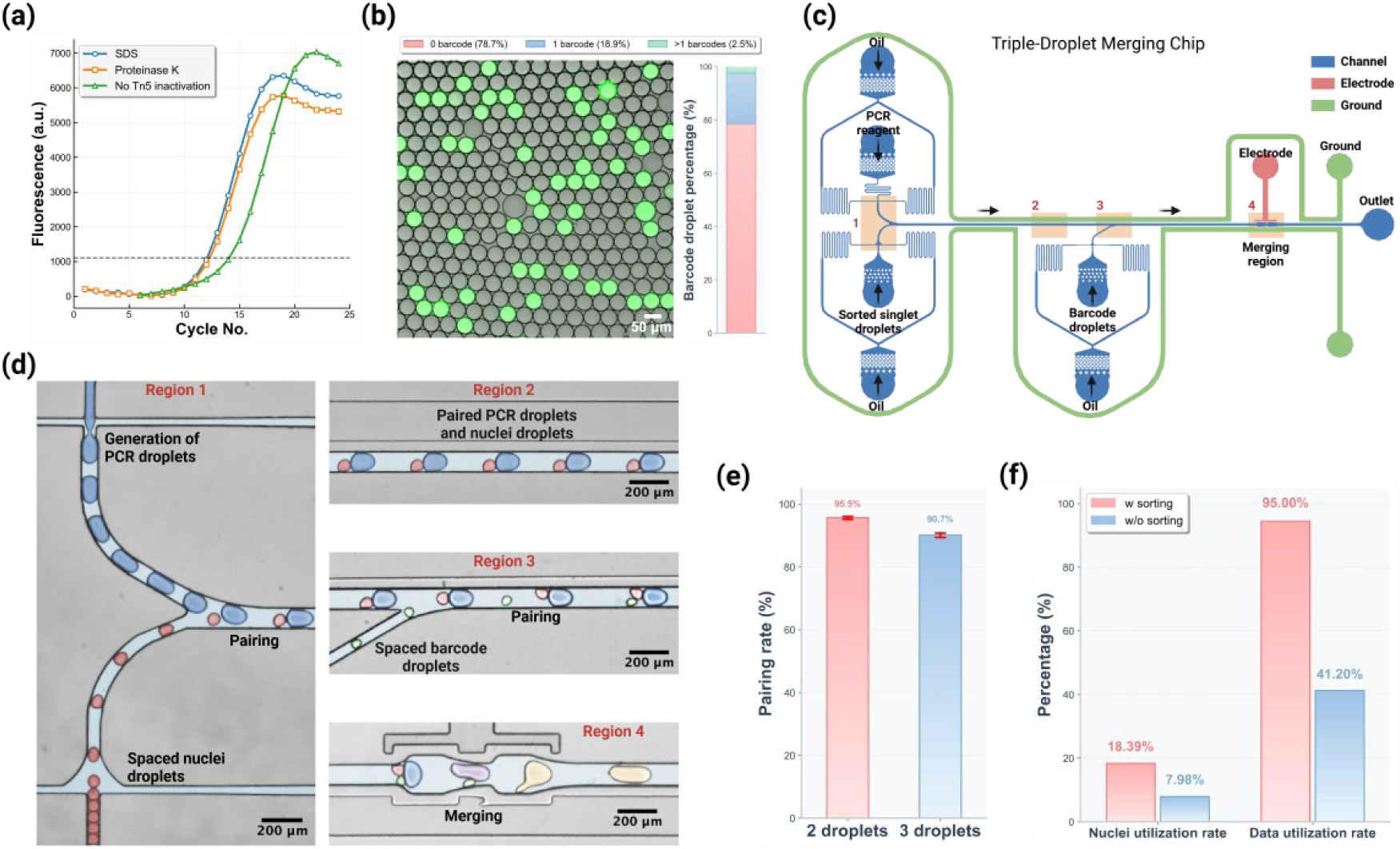
Validation of the Workflow of MusTer scATAC-seq. (a) qPCR results showing different amplification efficiency in Ct value from Tn5-tagged nuclei subjected to inactivation by SDS or proteinase K treatment compared to the control without Tn5 inactivation. (b) A representative microscopic image showing the barcode droplets, produced by ddPCR, where the fluorescently green droplets, stained by SYBR Green I, were considered positive, or those containing the barcodes. Quantitative analysis of the positive rate based on ∼3,000 droplets was then used to compute the barcode encapsulation rate shown in the bar graph, ∼18.9% and ∼2.5% containing single barcode template and multiple barcode templates, respectively. (c) Schematic of the triple-droplet merging chip layout, showing the labeled inlets for PCR reagents, oil, reinjected sorted nuclei droplet and as-prepared barcode droplets, as well as the outlet for merged droplets. Electrode channel was filled with 5 M NaCl solutions, and Ground channel was filled with 45 mM NaCl solutions. (d) Color-labeled microscopic images showing, (1) Region 1: On-chip generation of PCR droplet; (2) Region 2: Pairing of a PCR droplet and a reinjected singlet droplet; (3) Region 3: Matching of the PCR-singlet paired droplets and a barcode droplet; (4) Region 4: Merging of the triple-droplet triad. (e) The one-to-one pairing ratio of singlet droplets and PCR droplets, and matching rate of the triple-droplet triad. (f) Estimation of the nuclei utilization rate and data utilization rate with and without sorting. Error bars stand for mean ± standard variation with n = 3.

ddPCR was employed here to produce a barcode library for labeling the tagged nuclei open chromatin fragments using the barcode droplet generator (**Fig. S1b**)^20^. Briefly, droplets (v ∼ 0.07 nL) were generated by co-encapsulating the PCR reagents containing diluted double-stranded barcode templates (λ = 0.24), shown in Supplementary Table 1. About 90 µL droplets would be generated in 10 mins. The PCR cycle was optimized at 30 to produce approximately 1.07 × 10⁹ identical template copies per droplet. The positive rate of generated barcode droplets was characterized as 21.34% ± 1.45% (**Fig. 5b**). According to the empirically observed positive rate and Poisson statics^60^, approximately 19% of the barcoded droplets were estimated monoclonal, accounting for 90% of barcode template-containing droplets.

The triple-droplet merging chip (**Fig. S3**) was then introduced to fuse three droplets, *i.e.* PCR droplets, reinjected sorted nuclei singlet droplets and as-prepared barcode droplets. Noted that the PCR droplets contained the PCR buffer, reverse primers, dNTPs, and enzymes for in-droplet amplification after droplets merging. As shown in **Fig. 5c** and **5d**, four key regions were integrated into the triple-droplet merging chip, and flow rates were optimized carefully to allow the following tasks accomplished on the chip: (1) generation of the PCR droplets (Region 1), (2) pairing of a PCR droplet and a reinjected singlet droplet (Region 2), (3) matching of the PCR-singlet paired droplets and a barcode droplet (Region 3), and (4) merging of the triple-droplet triad (Region 4). Quantified as shown in **Fig. 5e**, one-to-one pairing of singlet droplets and PCR droplets (**Supplementary Movie 6**) and matching of the triple-droplet triad (**Supplementary Movie 7**) were characterized around 95.5% and 90.72%, respectively. The well-matched set of triple droplets was merged by DEP force *via* a non-uniform electric field generated by ionically conductive liquid electrodes (voltage of ±500 V at 50 kHz, **Supplementary Movie 8**).

Nuclei utilization rate, defined as the percentage of valid singlet droplets (single nucleus tagged with an appropriate barcode) relative to initial input nuclei numbers was characterized as 18.39% and 7.98%, equivalent to a processing speed of 3,148 and 1,365 singlet droplets/minute, with and without the multi-parametric selection (**Fig. 5f** and **Fig. S11**). The improved nuclei utilization rate is attributed to the ability of multi-parametric selection effectively isolating the singlet droplets. The nuclei utilization rate is ultimately lower than 20% owing to that the low positive rate of barcode droplets once again originated from the request of limited dilution. Nevertheless, this rate can be readily improvable by adopting to other barcoding methods, such as those using hydrogel beads.

Data utilization rate, defined as the percentage of singlet data that can be effectively analyzed post-sequencing relative to the total number of droplets containing nuclei reinjected into the triple-droplet merging chip, was around 95% with droplets sorting, more than twice of the counterpart without droplets sorting (**Fig. 5f**), suggesting that the sequencing efforts were made optimally utilized by the proposed MusTer scATAC-seq platform.

### Improved scATAC-seq of C4 Maize Leaf by MusTer

A key trade-off exists between throughput and data quality in scATAC-seq. The high-throughput 10X Chromium system^55^ fails to achieve the high-resolution chromatin accessibility profiles generated by the low-throughput PCR plate-based method^30, 56^. Applying MusTer to maize leaf, we break this trade-off by generating a high-throughput dataset that also achieves high data quality, outperforming both previous approaches (**Fig. 6**).

**Fig. 6.**
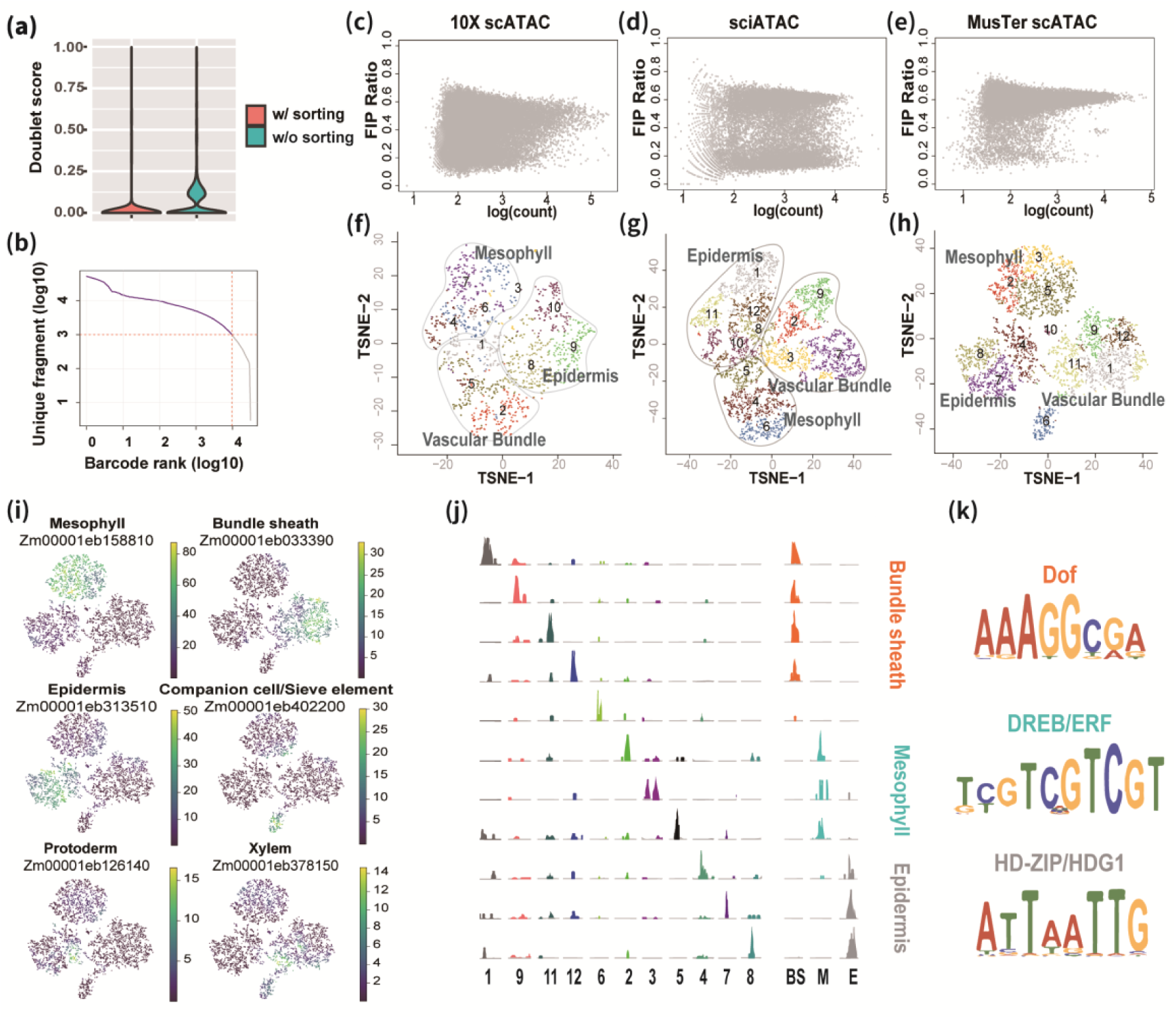
MusTer scATAC-seq for maize B73 leaves. (a) Doublet score of scATAC-seq using two Tn5 adapters with and without droplet sorting. (b) The number of unique Tn5 intergration sites per cell barcode. Fragment in promoter ratio per cell barcode for 10X scATAC-seq (c), sciATAC-seq (d) and MusTer scATAC-seq (e) data sets. t-SNE visualization of cell clustering for 10X scATAC-seq (f), sciATAC-seq (g) and MusTer scATAC-seq (h) data sets. (i) Maker genes annotation of clusters. (j) Genome track of identified top enriched accessible region for each cluster. (k) Enriched motifs discovered in three major clusters.

For example, MusTer’s on-chip multi-parametric singlet droplet sorting module reduced the raw averaged doublet events by almost five-fold (doublet rate of 23.3% versus 4.8%, **Fig. 6a**). After removing cells with less than 1,000 unique fragments, the doublet rate could be further reduced to 1.55%, compared to 4-10% doublet rate of 10X Chromium^24, 39^ and 6.0% of HyDrop^47^, showing effective nuclei aggregates removal by MusTer. Sequencing the library to ∼50% duplication rate and subsequent selecting for nuclei possessing ≥1,000 unique fragments yielded a final dataset of 9,120 nuclei, which had a median of 3,100 unique Tn5 integration sites per nucleus (**Fig. 6b**). Owing to the triple-droplet merging module, which facilitated more efficient Tn5 inactivation and optimal PCR conditions, the data enrichment achieved was significantly superior to the 10X system^55^. The quality metrics showed that around 90% of nuclei exhibit >45% of fragments in TSS (transcription start site) regions, which is much higher than ones with the 10X system^55^ (>30%) across all the read depth, suggesting a 50% increase in the signal-to-noise ratio (**Fig. S12**). Moreover, these nuclei showed >80% in accessible chromatin peaks that closely match or exceed those obtained from bulk ATAC-seq^67, 68^ (**Fig. S12**). FIP (fragment in promoter) ratio was also calculated using SnapATAC^40^ and compared with maize leaf scATAC-seq studies using 10X system^55^ and low throughput method (single cell indexed ATAC-seq, sciATAC-seq)^56^ respectively (**Fig. 6c, d**) using the same parameters. The result clearly demonstrated that our method yielded fewer cell barcodes with a low FIP ratio (<40%) (**Fig. 6e**). Cell barcodes with a FIP ratio below 40% are likely a result of background noise from empty droplets and damaged nuclei which can both be mitigated with MusTer. Overall, our scATAC-seq data indicated MusTer platform can be effectively adopted for scATAC-seq with high quality cells and low level of background signals. Next, we grouped the cells into three major clusters, which were further divided into 12 smaller subclusters using SnapATAC (**Fig. 6h**). Our data revealed sharper separation with well-defined boundaries and little undetermined cells compared with 10X system^55^ and PCR plate-based sciATAC-seq method^56^ (**Fig. 6f, g**). In addition, the two biological replicates are closely grouped in the cell clustering analysis (**Fig. S13**).

Annotation with established marker genes indicated that the three major clusters correspond to three primary cell types in leaves: mesophyll, epidermis, and vascular bundles (Supplementary Table 2). For instance, bundle sheath cells within the vascular bundle cluster were recognized by the well-characterized C4 marker gene ZmDCT2 (Zm00001eb033390), which plays a role in transporting the 4-carbon compound malate to the chloroplasts of bundle sheath cells for carbon fixation (**Fig. 6i**)^69^. Additional gene annotation with known markers also revealed the presence of xylem and companion cells within the vascular bundle cell cluster (**Fig. 6i**).

After peak calling using MACS2 for each of the 12 clusters, a total of 136,875 accessible chromatin regions were identified covering ∼3.3% of the maize genome. Differential accessible chromatin regions (dACRs) were also identified (**Fig. 6j**). In total, 3,041, 4,144, and 4,750 dACRs were found in mesophyll, bundle sheath and epidermis, respectively (Supplementary Table 3, 4, 5). Gene Ontology (GO) enrichment analysis of genes near the differentially accessible regions (dACRs) further confirmed the identity and function of each cell type (Supplementary Table 6, 7, 8). For instance, bundle sheath cells were enriched in pathways related to photosynthesis and carbon fixation (Supplementary Table 7). In contrast, epidermis was enriched in pathways associated with defense mechanisms and wax biosynthesis (Supplementary Table 8). Using a *de novo* motif discovery method, we identified the enriched DNA motifs for mesophyll, bundle sheath, and epidermis. For example, the dACR of the bundle sheath were found to be enriched with the Dof transcription factor motif, while those of the epidermis were enriched with the HDG1 motif, confirming that MusTer can accurately pinpoint potential master regulator in each cell type^56, 70^ (**Fig. 6k**).

## Discussion

The MusTer workflow provides a generalizable platform for droplet-based single-cell analysis by integrating singlet droplet sorting with multi-droplet merging strategies, thereby improving data quality, optimally utilizing sequencing capacity. Using maize scATAC-seq as a representative application, we have demonstrated that MusTer can effectively enrich the population of singlet droplets through a multi-parametric sorting strategy, achieving singlet purities above 95% even for nuclei samples that exhibits severe aggregation, performance that has not been achieved by existing methods. Though FADS has been successfully applied to high-throughput enrichment of rare events in enzyme evolution and antibody discovery^71–74^, these applications typically rely on a single sorting parameter, most commonly fluorescence intensity. Such a criterion is sufficient when fluorescence profiles are relatively simple. In contrast, singlet droplet discrimination requires higher-dimensional information. This limitation is illustrated in spinDrop^36^, which relies primarily on intensity-based gating and achieved only 77.2% singlet purity at λ = 0.5 in a cultured cancer cell line sample that typically shows minimal aggregation. MusTer overcomes this limitation by jointly exploiting fluorescence peak amplitude, width, and interval, enabling robust discrimination between singlet droplets and droplets containing doublets or small aggregates, even when fluorescence intensity differences are subtle. While extraction of peak shape features has been reported previously^75, 76^, to our knowledge, MusTer represents the first demonstration of their integration into a multi-parametric sorting framework for effective singlet droplet enrichment. In addition to outperforming existing droplet-based sorting strategies, multi-parametric droplet sorting offers several practical advantages over FACS-based singlet enrichment workflows. First, although FACS can isolate single cells or nuclei, subsequent droplet encapsulation still relies on limited dilution to minimize multiplet droplet formation, resulting in a large fraction of empty droplets and reduced effective throughput. In contrast, droplet-level sorting enriches singlet droplets directly, enabling a substantial higher proportion of nuclei to be processed productively. Second, FACS is typically operated at high flow rates, which can lead to considerable sample loss during sorting; for example, ∼50,000 input cells may be required to recover ∼1,536 sorted singlets^31^. In comparison, the multi-parametric sorting module in MusTer recovered ∼10,000 singlet droplets from ∼38,000 input nuclei, demonstrating markedly improved sample utilization. Finally, multi-parametric droplet sorting can be seamlessly integrated with the droplet encapsulation process, eliminating the need for separate upstream sorting steps and simplifying the overall experimental workflow (**Supplementary Movie 9**). Consistent with this multi-parametric droplet sorting capability, sequencing results show that the doublet rate was reduced from 23.3% to 4.8%, substantially mitigating doublet-associated artifacts and maximizing the effective use of sequencing reads. Moreover, cell barcodes derived from MusTer-processed samples exhibited a lower fraction of fragments in promoter (FIP) regions, an indicator of background noise, remaining below 40%, which is improved relative to prior scATAC-seq studies^54, 55^. Although effective, the current multi-parametric strategy for singlet droplet sorting still depends on careful tuning of experimental parameters, including PMT gain, droplet flow rate, and nuclear staining, to obtain well-separated clusters in the amplitude-width scatter plots, as well as on empirically defined gating criteria. Further development will therefore be required to improve the robustness, generalizability, and standardization of this approach across different samples and experimental conditions.

On the other hand, MusTer further enhances droplet-based scATAC-seq data quality by enabling key reaction steps to be performed under optimal conditions through droplet merging. The multi-droplet merging module allows precise matching and merging of three droplets containing distinct components, thereby decoupling otherwise incompatible biochemical steps. In the context of scATAC-seq, this capability enables the introduction of barcodes and PCR reagents only after effective denaturation of Tn5 transposase using proteinase K. In conventional one-pot droplet reactions, proteinase K treatment required for Tn5 inactivation would also inactivate PCR polymerase, leading to workflow failure. Denaturation of Tn5 transposase has previously been shown to improve data quality in plate-based scATAC-seq^34, 65, 66^; however, to our knowledge, MusTer represents the first demonstration of effective Tn5 denaturation in a droplet-based scATAC-seq workflow. Consistent with this design, MusTer scATAC-seq achieves a substantially higher fraction of fragments nearby TSS (45%), compared with data generated using the commercially available 10X Chromium system (∼30%)^55^. Moreover, when analyzed using the same processing and visualization pipeline reported previously for maize scATAC-seq^55, 56^, MusTer-derived data exhibit sharper cluster separation and more clearly defined boundaries among the three major cell populations, indicating improved signal-to-noise and biological resolution. Beyond scATAC-seq, the multi-droplet merging strategy is considered a generalizable strategy for improving data quality in droplet-based single-cell workflows that require complex, multi-steps reactions. Unlike pico-injection, which is limited to the addition of homogeneous solutions, multi-droplet merging enables the modular combination of droplets containing distinct reagents or barcoded entities. For instance, in workflows involving cell lysis and reverse transcription (RT)^19^ or PCR within droplets, lysis can be performed independently under optimal conditions before merging with droplets containing enzymatic reagents and barcode templates, thereby improving reaction efficiency and flexibility. Despite these advantages, MusTer still faces technical challenges, including potential perturbations to droplet stability during the reinjection process. This limitation may be mitigated by further integrating nuclei droplet generation and singlet droplet sorting on a single microfluidic platform, as demonstrated in **Supplementary Movie 9**.

## Materials and Methods

### Materials

Oligos used in this study were designed as shown in Supplementary Table 1 (synthesized and purified by Sangon Biotech Vendor).

### Design and Fabrication of Microfluidic Chips

The patterns of microfluidic chips (Fig. S1, **2, 3**) used in this study were designed using CAD software (Autodesk, San Rafael, USA) and subsequently printed on film or chromium/gold (Cr/Au) photomasks (MicroCAD Photo-Mask Ltd., Shenzhen, China). The chips were fabricated via standard soft-lithography process, replicating elastomers polydimethylsiloxane (PDMS) on the SU-8 mold, followed by bonding on glass slides to seal the channels.

Specially, the SU8 mold for the nuclei droplet generator (**Fig. S1a**) was fabricated on a silicon wafer pre-cleaned by acetone and isopropanol. SU8-3050 photoresist (Kayaku Advanced Materials, USA) was spin-coated on the silicon wafer and pre-exposure baked (95°C, 15 min). Through exposure photomask with proper dosage (365-nm UV, ABM Mask Aligner, USA), the patterns were transferred onto the silicon wafer. After post-exposure bake (65°C for1 min followed by 95°C for 5 min), the uncross-linked photoresist was removed by the SU8 developer (MicroChem, USA) (5 min agitation), followed by a hard bake (110°C, 15 min) to strengthen the structures. The fabricated mold was measured approximately 45 µm in height. The mold for the barcode droplet generator (**Fig. S1b**) and the triple-droplet merging chip (**Fig. S3**) were fabricated with a height of 37 μm and 69 μm, respectively.

PDMS base and curing agent (Sylgard 184, Dow Corning, USA) were mixed thoroughly with a weight ratio of 10 : 1 and poured onto the fabricated SU-8 master molds. After degassing for 30 minutes and curing at 70°C for 2 hours, the cured PDMS were peeled off from the molds and cut into slabs. Holes for inlets and outlets were created with a 1.0 mm Miltex biopsy puncher (Integra Life Sciences, USA). The PDMS slabs were subsequently bonded with cover glasses spined-coated by a layer of semi-cured PDMS (weight ratio of base to curing agent set at 2 : 1 and semi-cured at 110°C for 1 min). The assembled microfluidic chips were then baked at 80℃ for 12 hours in an oven to strengthen the bonding.

The SU8 mold for the droplet sorting chip (**Fig. S2**) was fabricated 40 μm in height. PDMS replicas fabricated and hole-punched similarly as above-described were boned with the non-conductive side of an ITO-coated glass slides through PDMS bonding^75^. Electrode was then integrated employing a vacuum-assisted molten alloy filling process. Briefly, the bonded chip was heated to 110°C on a hotplate to melt solder alloy wire (70°C melting point) within electrode channels. Negative pressure applied through outlet tubing enabled complete channel filling, after which contact pins were inserted into the inlets. Cooling the alloy to room temperature established stable electrical contacts between contact pins and on-chip electrodes verified by a multimeter.

### The Multi-Parametric Singlet Droplet Sorting System

The multi-parametric singlet droplet sorting system was custom-developed based on previously reported architectures with substantial hardware modifications (**Fig. S4**)^75, 77^. This system integrated multiple functions, including bright-field imaging, laser-induced fluorescence detection, real-time signal processing and sorting actuation. Bright-field illumination was provided by a white light LED coupled with a 676/29 nm bandpass filter (86-988, Edmund Optics Inc., USA), and images were captured by a high-speed CMOS camera (EoSens 2.0 MCX12-FM, SVS-Vistek GmbH, Germany). Fluorescence excitation of the flowing droplets was achieved using a 488-nm laser (MDL-III-488L-50mW, CNI Optoelectronics Tech. Co. Ltd., China), focused through a 10x objective (MRH20101, Nikon Instruments Inc., Japan). The laser beam (output near TEM00 mode, M^2^ factor < 2.0) was expanded in one dimension by 2 cylindrical lenses to form a rectangular shape (**Fig. S5**), with its major axis oriented perpendicular to the channel and fully covering the 30 µm channel width, thereby enabling the detection of internal features within the droplets. The emitted green fluorescence was detected by a photomultiplier tube (PMT, PMM02, Thorlabs Inc, USA), generating an analog voltage signal subsequently converted to digital data by a 12 bit analog-to-digital converter (ADC, AN9238, Alinx Electronic Ltd., China) at a sampling rate of 1 MSPS. Real-time signal processing was performed on a hybrid FPGA/CPU architecture (AX7Z100, Alinx Electronic Ltd., China), where the Programmable Logic (PL) core handled data analysis and sorting trigger generation at a clock rate of 1 MHz, while the Processing Subsystem (PS) core managed communication with the host PC through Ethernet. A signal generator (SDG 1022X, SIGLENT Technologies Co. Ltd., China) received sorting trigger to produce square waves, which are then amplified to several hundred volts by a custom-built voltage amplifier. These high-voltage pulses created a non-uniform electric field on-chip (**Fig. S14**), enabling dielectrophoretic (DEP) force-mediated droplet sorting.

### Preparation of Droplets Encapsulating Fluorescent Microbeads

FITC-labeled polystyrene (PS) microbeads (4-μm and 10-μm, KBsphere^®^, Suzhou Knowledge & Benefit Sphere Tech. Co., Ltd., China) were suspended in an aqueous solution containing 20% OptiPrep^TM^ (D1556, Sigma-Aldrich, Germany) and 0.1% (w/w) Tween-20 (P1379, Sigma-Aldrich, Germany). OptiPrep^TM^ was added to increase the density of the suspension and minimize sedimentation of the PS beads during droplet generation, while Tween-20 served to reduce bead aggregation by acting as a surfactant. The suspension was briefly sonicated prior to use to further disrupt potential aggregates. The concentration of bead suspension was adjusted according to the intended number of beads encapsulated per droplet. Droplets were generated using the nuclei droplet generator (**Fig. S1a**). The prepared aqueous bead suspension was introduced onto the chip at a flow rate of 4.9 μL/min, while the continuous oil phase (Droplet Generation Oil for EvaGreen, Bio-Rad Laboratories Inc., USA) was delivered at 21 μL/min. Under these conditions, monodisperse water-in-oil droplets with an average diameter of ∼60 μm were generated and collected into microcentrifuge tubes. For experiments requiring a mixed droplet population containing both 4-μm and 10-μm beads, the two types of droplets were generated separately and then combined at a defined volumetric ratio. Gentle pipette agitation was applied to ensure homogeneous mixing of the two droplet populations without inducing droplet coalescence.

### Growth of Maize, Sampling and Nuclei Isolation

Maize seeds (*Zea mays* B73) were grown under the conditions of 12:12 Light/Dark cycles at 30 °C Light/22 °C Dark and at 50% humidity. Leaves were collected from 10-day old maize seedlings and chopped into small pieces in nuclei isolation buffer (Supplementary Table 9) to release nuclei. 0.2% (v/v) of formaldehyde (F1635, Sigma-Aldrich, Germany) was then added to fix nuclei on ice for 5 minutes, followed by addition of 0.2% (v/v) Triton X-100 (cat. no. A16046.A, Invitrogen, US) for nuclei membrane permeabilization and chloroplast lysis. The solution was then filtered through a 40 μm cell strainer (CLS431751, Corning-Sigma-Aldrich, Germany) and nuclei were pelleted by centrifugation at 500g, 4°C for 10 min. Then the pellet was resuspended with nuclei isolation buffer and filtered through a 10 μm nylon filter (NY1004700, Sigma-Aldrich, Germany). The nuclei were pelleted again by centrifugation and washed one more time with the tagmentation wash buffer (Supplementary Table 9). Before the final centrifugation, a small fraction of the solution was removed and stained with SYBR Green I (10000x, Thermo Fisher, USA) to determine the nuclei concentration.

### Tn5 Transposase Tagmentation and Nuclei Droplets Generation

The expression and purification of TS-Tn5 tagmentation enzyme were conducted using vector obtained from Addgene (accession number 127916) and following the method described previously^78^. Two sets of Tn5 adapter oligos with distinct barcode sequences (Supplementary Table 1) were used in the Tn5 transposition process for the post-sequencing assessment of nuclei clumps *i.e.* doublets. Pre-annealed Tn5-ME-A and Tn5-ME-B adapters (50 μM, supplementary Table 1) were mixed 1:4 with TS-Tn5 proteins (1 OD/ul) and incubated at 25°C for 1 hr to assemble the Tn5 transposon. About 1.2 × 10^5^ nuclei were suspended in 250 μL tagmentation buffer (Supplementary Table 9) with 10 OD assembled transposon. The tagmentation reaction was carried out for 45 minutes at 37°C with rotation. The tagmentation reaction was stopped by adding 1 mL nuclei wash buffer (Supplementary Table 9) supplemented with 10 mM EDTA (15575020, Thermo Fisher, USA) to the solution. The nuclei were pelleted by a centrifugation at 500g, 4°C for 10 min. The nuclei were then washed with nuclei wash buffer twice and filtered through 10 μm nylon filter. The nuclei were finally resuspended in nuclei wash buffer and the nuclei concentration were adjusted to around 2,000 nuclei/μL.

For tagmentation with two different Tn5 adapters, the nuclei were split into two parts and mixed with two different transposons to carry out tagmentation separately. The nuclei tagged with two different Tn5 adapters were then washed with nuclei wash buffer twice and filtered through 10 μm nylon filter separately. After nuclei counting, same number of tagged nuclei with two different Tn5 adapters were mixed in nuclei buffer (Supplementary Table 9) prior to the step of droplets encapsulation.

Nuclei droplets were subsequently generated after Tn5 transposition and 5× SYBR Green I staining using the nuclei droplet generator (**Fig. S1a**). Two aqueous phases, tagged nuclei in nuclei buffer supplemented with 14% (v/v) OptiPrep and 2% (v/v) thermolabile proteinase K (P8111L, NEB, USA) in nuclei buffer, were emulsified by the Bio-Rad oil phase (**Fig. S15a**). The flow rate for tagged nuclei, proteinase K and oil was 4.9 μL/min, 4.9 μL/min and 21 μL/min respectively for stable droplets generation with intended size.

### Barcode Droplets Generation

Prior to generating barcode droplets, double-stranded barcode templates were generated by hybridizing the two strands (maintaining the structural stability and thus consistent positive rate after droplet PCR. About 30 ng of single-stranded barcode templates were added to a 50 μL PCR reaction mixture, followed by 5 cycles of PCR amplification using KAPA HiFi HotStart polymerase (KK2502, Roche, Switzerland). After amplification, the products were purified with a PCR purification kit (MinElute PCR Purification Kit, QIAGEN, Germany), and the concentration was measured using a fluorometer (Quantus Fluorometer, Promega, USA). Barcode droplet generator (**Fig. S1b**) was utilized to produce 90 μL barcode droplets containing PCR reagents with 10 μL/min aqueous flow rate and 17 μL/min oil flow rate, which included 0.02 pg double-stranded barcode templates, 0.2 μM forward and reverse primers, 1x PCR buffer, 0.4% (v/v) Tween-20, 0.4% (m/v) polyethylene glycol (PEG) 8000 (Promega, USA), and KAPA HiFi polymerase (KK2502, Roche, Switzerland). Following the generation of barcode droplets (**Fig. S15b**), 35 μL of them were transferred into each PCR tube, and 35 μL of mineral oil was added to each tube to prevent droplets evaporation and coalescence during amplification. The droplet PCR was carried out under the following thermal cycling conditions: 3 minutes at 95℃; 30 cycles of 15 seconds at 95℃, 15 seconds at 63℃, and 15 seconds at 72 ℃; followed by a final extension of 2 minutes at 72℃, with all temperature ramp rates set to 0.8℃/s. To evaluate the positive rate, 20 μL of droplets were taken for an additional 15 PCR cycles and then stained with SYBR Green I for 20 minutes. The amplified products were verified by breaking the droplets to obtain the products, followed by agarose gel electrophoresis (**Fig. S16a**).

### Proteinase K Activation and Triple-Droplets Merging

After the sorting of nuclei singlet droplets with proposed multi-parametric selection method, the selected droplets encapsulating nuclei singlets were transferred into a PCR thermocycler and covered with mineral oil. The nuclei droplets were initially incubated at 37°C for 20 minutes to denature Tn5 proteins by thermolabile proteinase K activation, followed by incubation at 55°C for 15 minutes to inactivate proteinase K. The triple-droplet merging chip was firstly prefilled with 5 M sodium chloride solution (NaCl, S9888, Sigma-Aldrich, Germany) in the ground channel at 10 μL/min and 45 mM NaCl in the electrode channel at 0.9 μL/min to prepare aqueous salt bridge electrodes. After producing barcode droplets with a set positive rate, they were reinjected into the triple-droplet merging chip along with 300 μL of HFE-7500 oil (3M™ Novec™, USA) placed at the bottom of a 1 mL syringe (Becton, Dickinson and Company, USA). For small volumes of sorted singlet droplets, 300 μL of HFE-7500 oil was added at the bottom of the 1 mL syringe before merging to ensure complete expulsion of all sorted singlet droplets. PCR droplets were generated in real-time on the triple-droplet merging chip encapsulating PCR reagents including KAPA buffer (KK2102, Roche, Switzerland), and KAPA polymerase, dNTPs, reverse primers, Tween-20, PEG 8000 and BSA (Thermo Fisher, USA). To optimize the accuracy in two-droplet and three-droplet pairing, as well as to reduce the probability of multiple nuclei singlet droplets pairing with barcode droplets at the same time, the flow rates were set as follows: Bio-Rad oil (for separating reinjected droplets) at 10 μL/min, PCR droplets generation Bio-Rad oil at 7.2 μL/min, reinjected nuclei singlet droplets at 2.35 μL/min, reinjected barcode droplets at 1.38 μL/min, and KAPA PCR reagent at 7 μL/min. A signal generator (DG1022, RIGOL, China) and a high voltage amplifier (AMT-5B20-LC, Matsusada Precision, Japan) were used to provide square waves with voltage of ±500 V at 50 kHz to the on-chip electrode, achieving a merging efficiency of 100% for droplets in the merging area. The merged droplets were shown in **Fig. S15c**.

### Preparation of the Sequencing Library

Following the triple-droplet merging, the merged droplets were placed into PCR tubes with mineral oil on top to prevent droplets evaporation and coalescence. Droplet PCR was conducted as the following: 15 min at 72 ℃, 3 min at 95 ℃; 12 cycles of 20 s at 98 ℃, 30 s at 63 ℃, 50 s at 72 ℃; and a final step of 2 min at 72 ℃ with all ramp rates set to 0.8 ℃/s. After PCR amplification, mineral oil was removed and 50 µL TE buffer was added to the top of the emulsion followed by 30 µL of 100% droplet releasing agent *1H,1H,2H,2H*-perfluoro-1-octanol (Sigma-Aldrich, 370533). The samples were briefly vortexed and centrifuged, and the aqueous phase were purified using 0.9x AMPure XP beads (Beckman, USA). Subsequently, another round of PCR amplification using KAPA HiFi was carried out to add i5 and i7 sequencing adapters on the strand, producing the final sequencing library. The library size was determined by gel electrophoresis prior to sequencing (**Fig. S16b**).

### scATAC-seq Data Processing

The cell barcode sequences were extracted from the read1 fastq file to identify all the barcode sequences. Cell barcodes with reads number larger than 100 were kept and associated reads were used for further processing. The ratio of reads from the Tn5 barcodes was used to calculate a doublet score. The corresponding barcode sequence were added to the name of each read and the fastq file then further mapped to Zm-B73-REFERENCE-NAM-5.0 maize genome using bowtie2 to generate the BAM file. Quality control analysis was performed using Socrates R package^55^. The BAM file was processed using Snaptools to generate snap file as input data file for SnapATAC for cell clustering and annotation analysis^79^. Cell barcode with more than 1000 reads and FriP (fragment in promoter) ratio higher than 0.4 were kept. Bin size of 500bp was chosen and bins were then sorted and filtered to remove the top 5% invariant features. Dimension reduction was then performed, and the first five eigenvectors were used for the downstream clustering analysis. Clustering was visualized via t-SNE implemented by Rtsne. Two previous maize leaf scATAC-seq data (GEO Accession: GSM4696890, NCBI SRA: SRX23266769) were processed using SnapATAC with the same parameters. Accessible regions and differential accessible regions (DARs) for each cluster were identified using SnapATAC R package^40^. *De novo* motifs discovery for DARs were identified by using XSTREME (version 5.5.3) of the MEME suite (version 5.5.5) package^80^.

### Statistical Analysis

To quantify the percentage of each droplet type (singlet and multiplet droplets), fluorescence images were acquired for each droplet population under both pre- and post-sorting conditions. Manual counting was performed on multiple images for each droplet population. Within each droplet population, droplet counts for each type were aggregated across all acquired images, and percentages were calculated by normalizing to the total number of droplets containing beads or nuclei (excluding empty droplets).

Given that the acquired images represented only a small fraction of the total droplet populations, a bootstrap resampling approach was employed to provide robust error estimates. Bootstrap analysis was performed with 1,000 iterations, where images were randomly resampled with replacement from the original dataset for each iteration. The resulting bootstrap distribution was used to calculate 95% confidence intervals, which were subsequently used as error bars in data visualization.

## Acknowledgements

This work was supported by the Research Grants Council of Hong Kong Special Administrative Region, China (project #: CUHK 14219922, 14207424, 14109420, N_CUHK410/24 and C5005-23W). We also thank Jin Chen for the assistance on fabricating the triple-droplet merging chips.

## Author Contributions

L.L., W.L., G.C., S.Z. and Y.P.H. conceptualized the study. L.L. developed and optimized nuclei droplets, barcode droplets workflow. L.L. and F.Q. developed and optimized the droplet merging workflow. L.L. and W.L. optimized the single-cell ATAC-seq workflow and W.L. performed downstream sequencing data analysis. G.C. set up and optimized the droplet sorting platform and G.C., L.L. optimized the multi-parametric singlet droplet sorting workflow. L.L., G.C. and W.L. co-wrote the manuscript, with input from F.Q., S.Z. and Y.P.H. S.Z. and Y.P.H. supervised the work.

## Competing interests

The authors declare that they have no competing interests.

## Supplementary Information

**Fig. S1.**
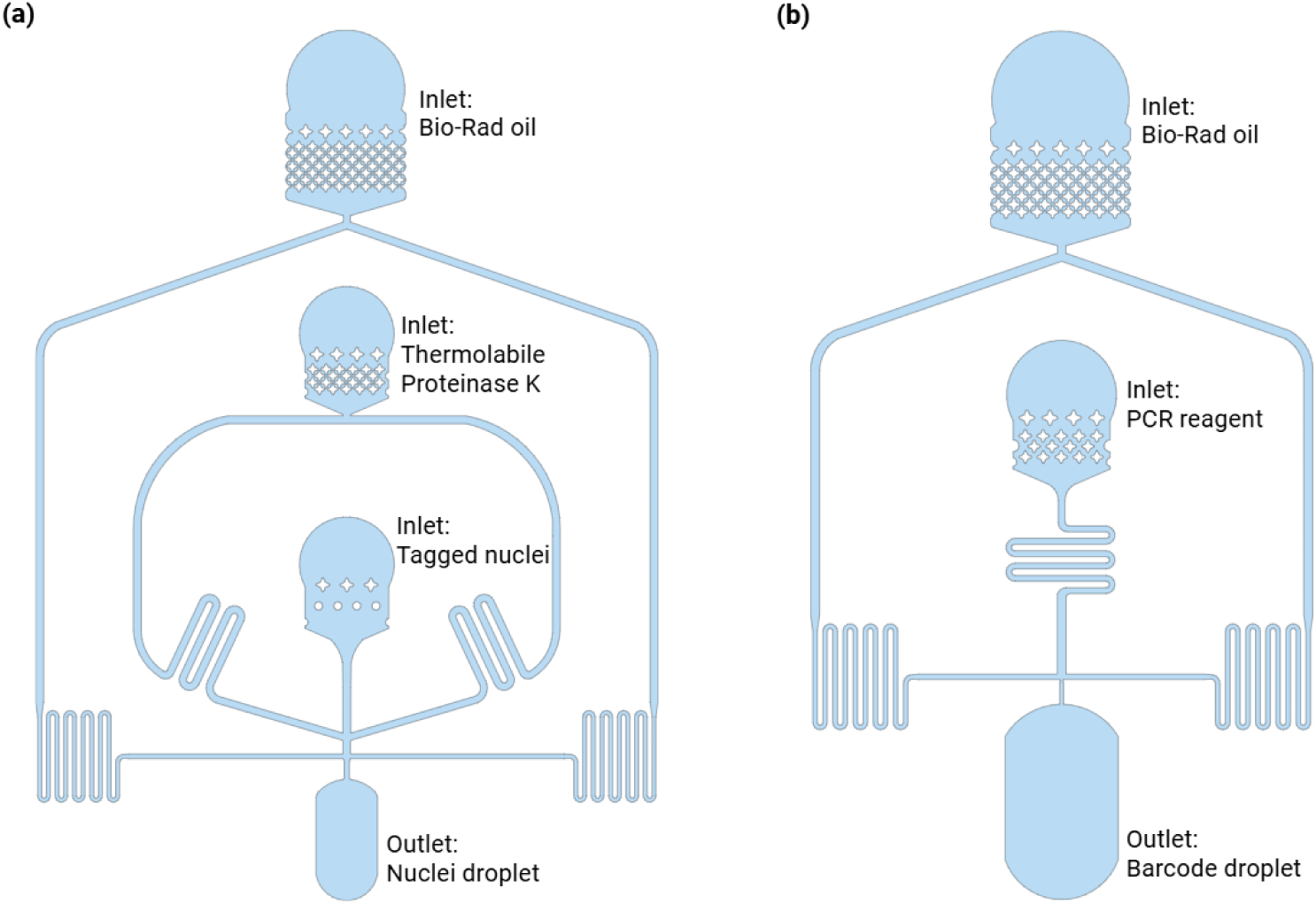
Design of (a) the Nuclei Droplet Generator and (b) the Barcode Droplet Generator.

**Fig. S2.**
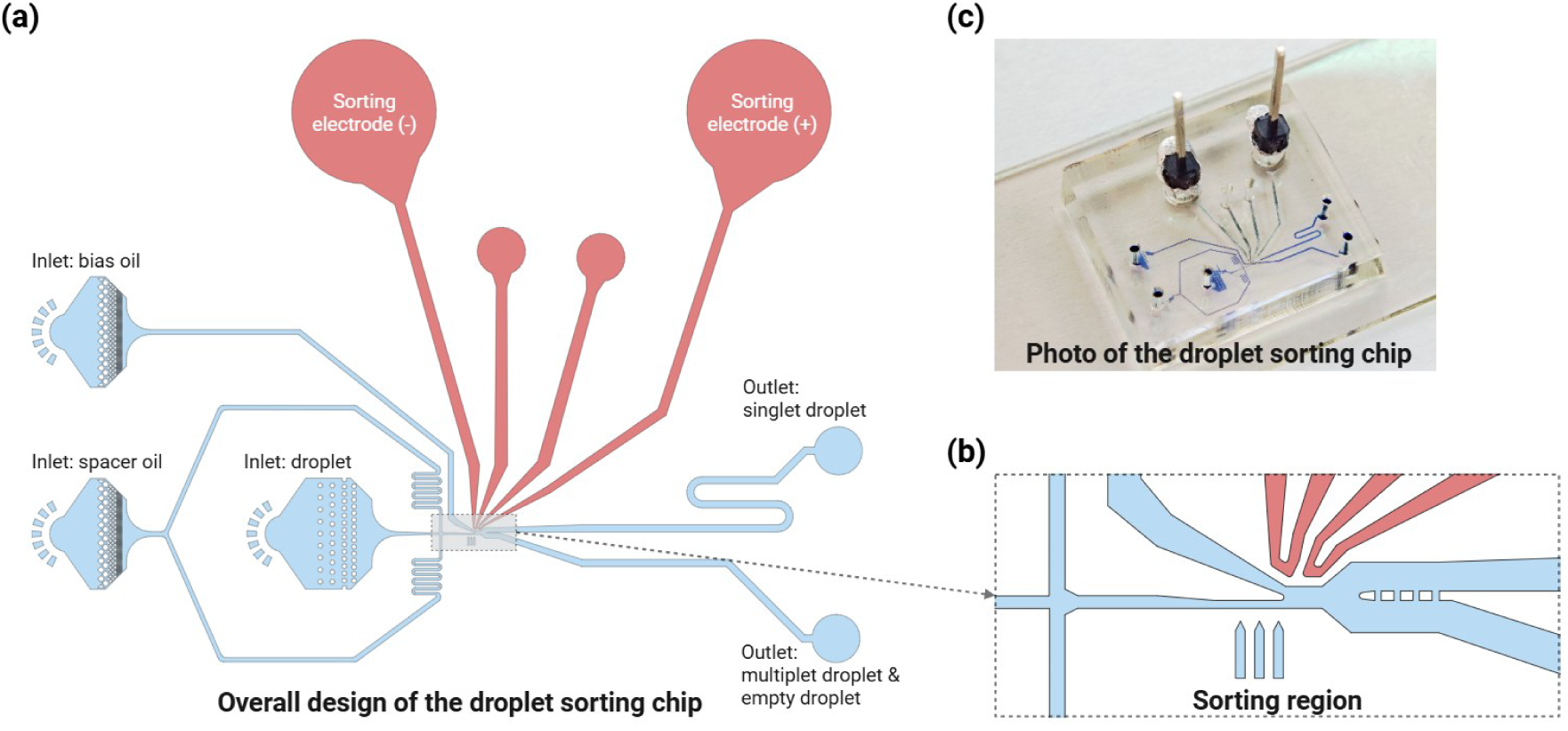
Design and Fabrication of the Droplet Sorting Chip. (a) Schematic of the design patterns, showing the inlets for droplets and oil, as well as the outlets for sorted singlet and unwanted multiplet or empty droplets. The spacer oil is used to separate the reinjected droplets, while the bias oil is used to push droplets into the outlet for multiplet and empty droplets in the absence of dielectrophoretic (DEP) force. (b) Enlarged view of the sorting region, where the singlet droplets are deflected by DEP force towards the designated outlet. (c) Photograph of the assembled droplet sorting chip inserted with the electrical connecting pins.

**Fig. S3.**
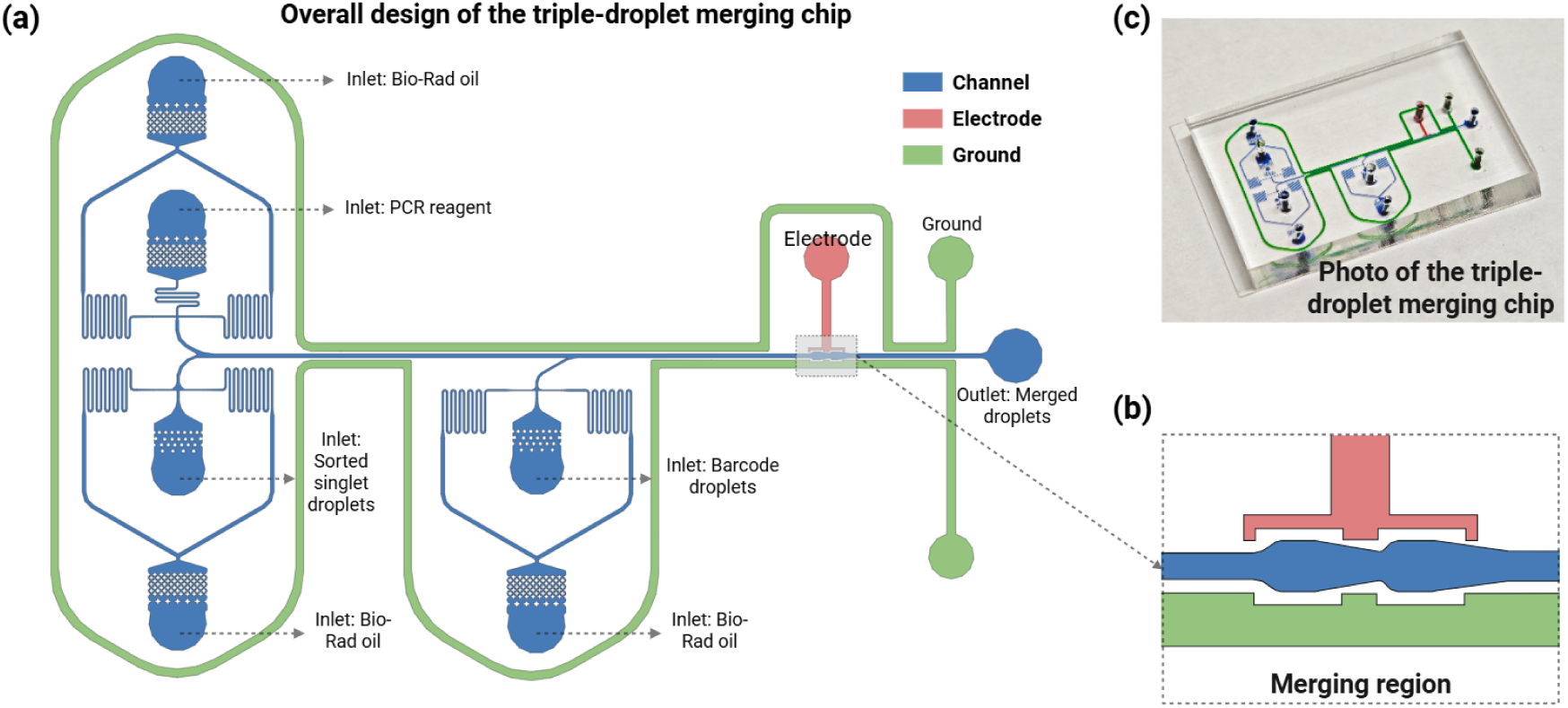
Design and Fabrication of the Triple-Droplet Merging Chip. (a) Schematic of the full chip layout, showing the labeled inlets for reinjected sorted singlet droplets, PCR reagents, as-prepared barcode droplets, salt solutions, and Bio-Rad oil, as well as the outlet for collecting the merged droplets. (b) Enlarged view of the merging region, where droplets are merged under an electric field. (c) Photograph of the actual fabricated triple-droplet merging chip.

**Fig. S4.**
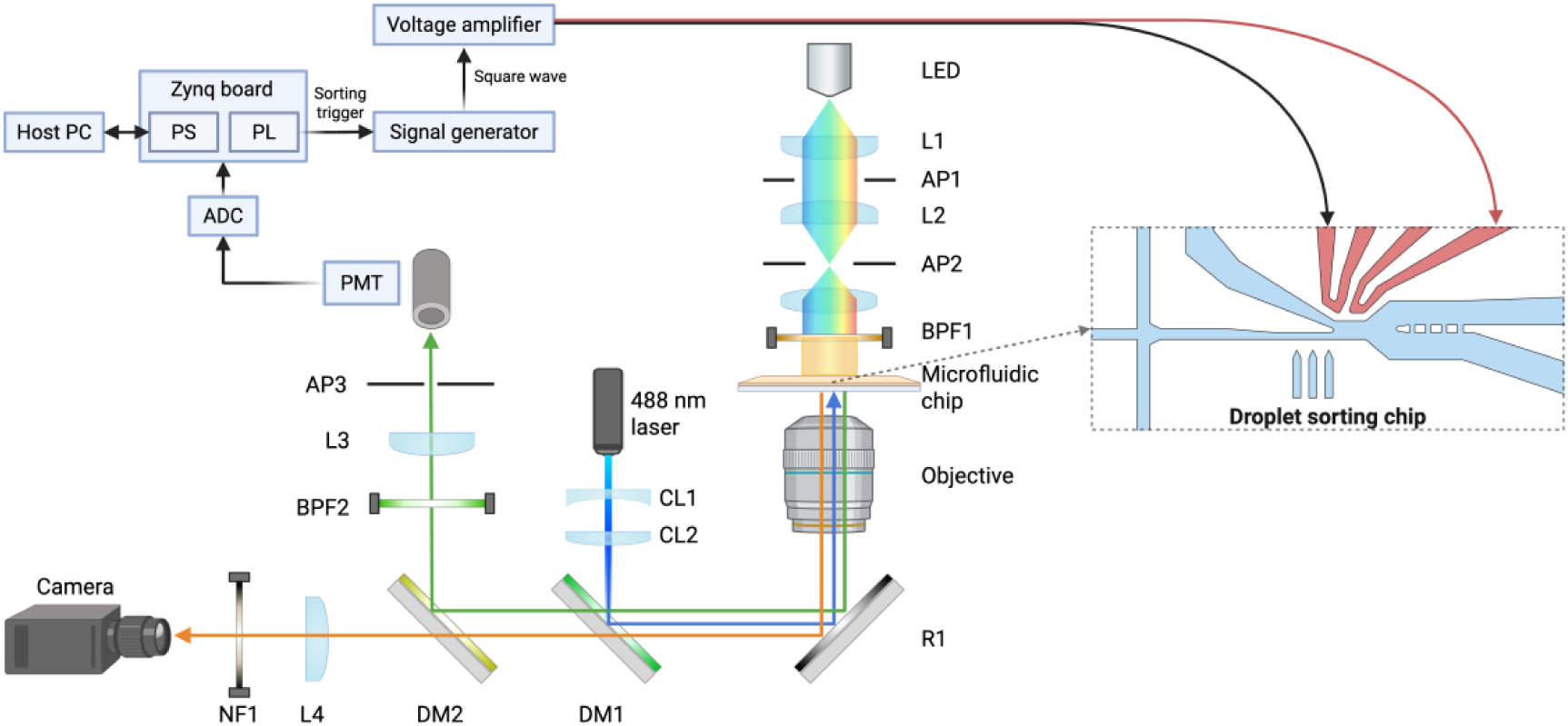
Schematics of the Multi-Parametric Singlet Droplet Sorting Module. The system integrates bright-field imaging, laser-induced fluorescence detection, real-time signal processing, and droplet sorting. A white LED with a 676/29 nm bandpass filter provides bright-field illumination, while a 488-nm laser, shaped by cylindrical lenses into a line focus, illuminates the flowing droplets. Emitted fluorescence is detected by a PMT and digitized by an ADC. Real-time peak analysis and sorting decisions are executed by a hybrid FPGA/CPU architecture (PL and PS), which communicates with a host PC. Sorting is triggered via a signal generator and voltage amplifier in response to FPGA output. Abbreviations: LED, light-emitting diode; L, lens; CL, cylindrical lens; AP, aperture; BPF, band-pass filter; R, reflector; DM, dichroic mirror; NF, notch filter; PMT, photomultiplier tube; ADC, analog-to-digital converter; PS, processing system; PL, programmable logic.

**Fig. S5.**
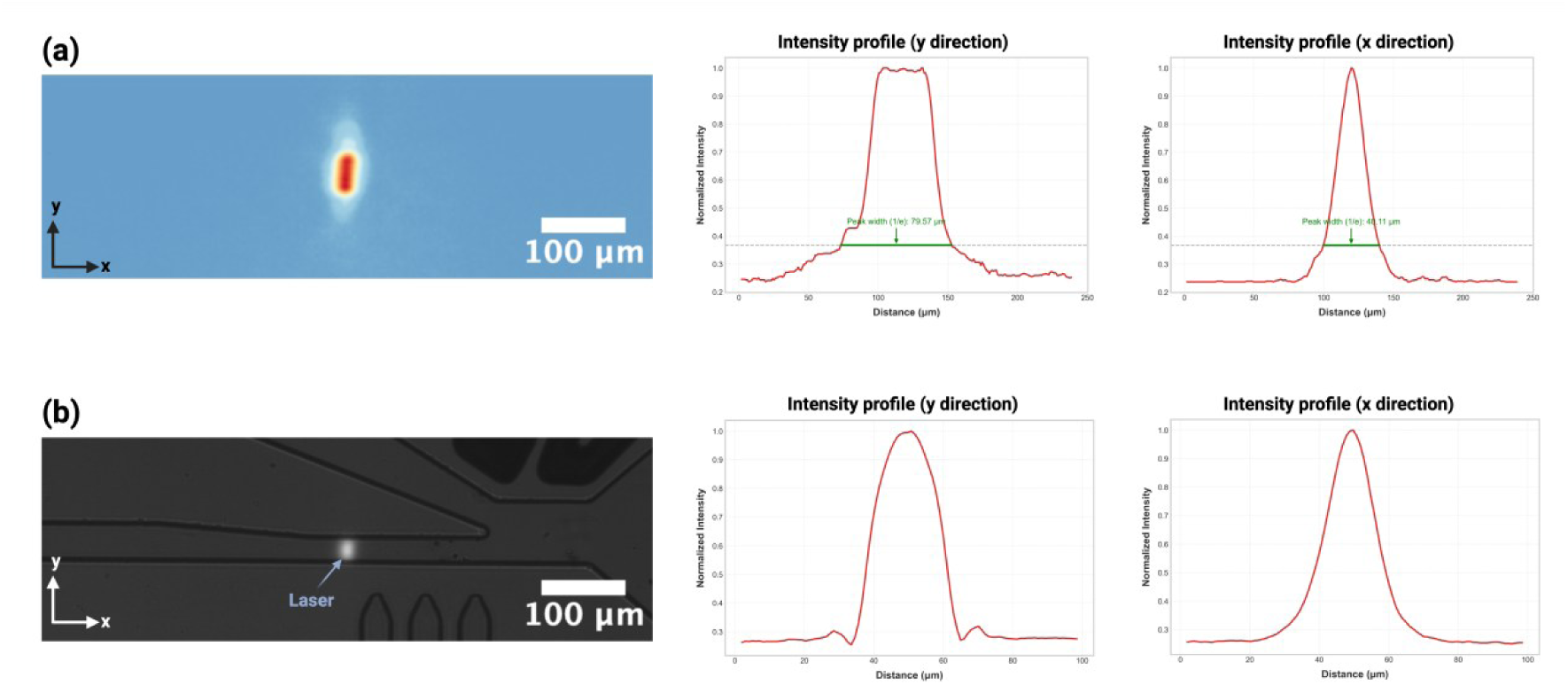
Visualization and Characterization of the Laser Beam Profile. (a) The laser beam profile was visualized by illuminating a fluorescein isothiocyanate (FITC, 0.05 mg/mL) solution with the fluorescence captured by a CMOS camera. The raw grayscale image was converted into a false-color spectrum to illustrate the spatial intensity distribution. Intensity profiles were extracted along both the x- and y-axes. The beam was configured as a rounded rectangle, with a major axis length of approximately 80 μm (along the y-axis) and a minor axis width of approximately 40 μm (along the x-axis), both defined at the 1/e maximum intensity threshold. Along the y-axis, the central 30 μm region displayed a relatively uniform intensity, which ensures consistent excitation as droplets traverse the detection region. (b) The laser beam was precisely aligned with the microfluidic channel such that its center coincided with the channel axis. The beam fully covered the 25 μm-wide channel, as visualized by the FITC fluorescence, ensuring complete excitation across the droplet path.

**Fig. S6.**
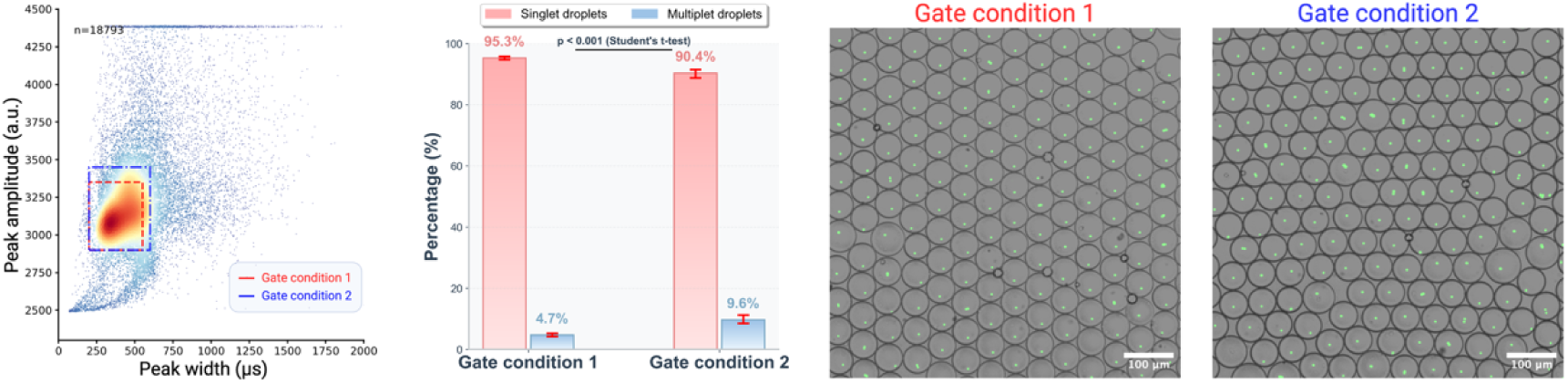
Gating Stringency on the Purity of Isolated Droplets. Droplets encapsulating fluorescent 4-μm PS beads, mimicking both singlets and multiplets encapsulated in drops, were sorted using gating of differing stringency. **Left**: Scatter plot of fluorescence peak amplitude versus peak width from 18,793 events, with the two different gated regions highlighted: Gate 1 (relatively strict, red dashed box) and Gate 2 (relatively liberal, blue dashed box). **Middle**: Quantification of singlet and multiplet droplet fractions post-sorting. Gate 1 yielded 95.3% singlet and 4.7% multiplet droplets, respectively. Under gate 2, the purity of singlet droplet was 90.4% whereas the multiplet population was 9.6% (p < 0.001, Student’s t-test). **Right**: Representative microscopic images of sorted droplets validated that stricter gating (Gate 1) effectively excludes multiplets, whereas more liberal gating (Gate 2) permited more multiplet-containing droplets. Error bar represented 95% confidence interval (droplet number N = 2,623 for gate condition 1 and N = 2,081 for gate condition 2).

**Fig. S7.**
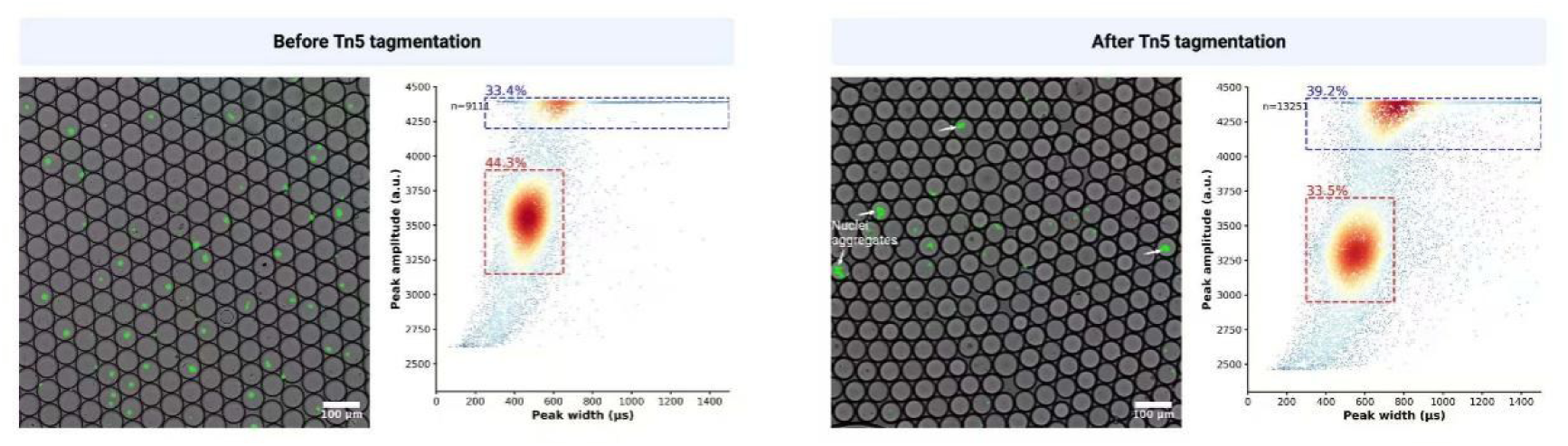
Tn5 Transposase Tagmentation on Nuclei Aggregation. **Left**: A representative microscopic image of droplets encapsulating nuclei before Tn5 transposase tagmentation and amplitude-width scatter plot from 9,111 peaks, with the singlet droplets cluster gated in the red dashed box and the multiplet droplets cluster gated in the blue dashed box. **Right**: A representative microscopic image of droplets encapsulating nuclei after Tn5 transposase tagmentation and amplitude-width scatter plot from 13,251 peaks, with the singlet droplet cluster gated in the red dashed box and the multiplet droplet cluster gated in the blue dashed box.

**Fig. S8.**
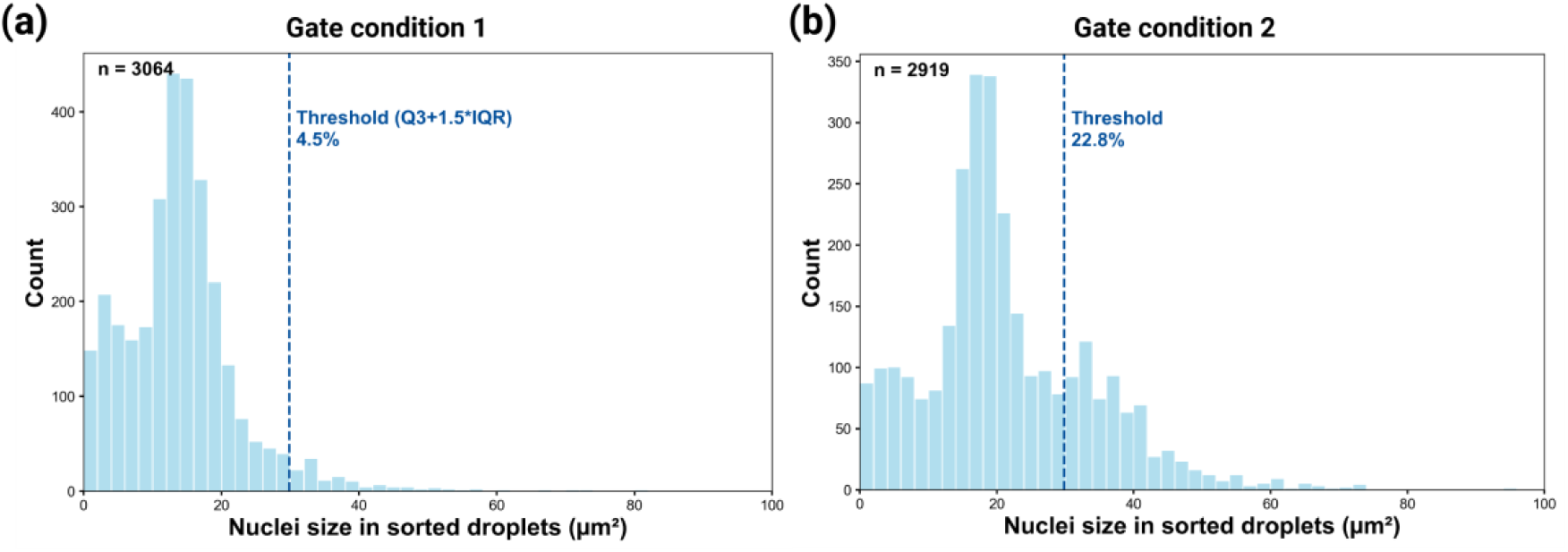
Size Distribution of Nuclei in Sorted Droplets Obtained Under Two Gate conditions. To quantify the proportion of multiplet droplets (Types III–IV) arising from nuclei aggregates, nuclei sizes were measured from post-sorting fluorescence images. (a) Under Gate condition 1, where the aggregate proportion was minimal, an upper size threshold was defined as 1.5 × the interquartile range above the third quartile (Q3+1.5*IQR) and used as a statistical cutoff for aggregate identification. Nuclei with sizes exceeding this threshold were classified as aggregates. Using this method, it was calculated that multiplet droplets (Type III - IV) accounted for 4.5% under Gate condition 1. (b) The same threshold was subsequently applied to droplets sorted under Gate condition 2, where the proportion of multiplet droplets (Type III-IV) was 22.8%.

**Fig. S9.**
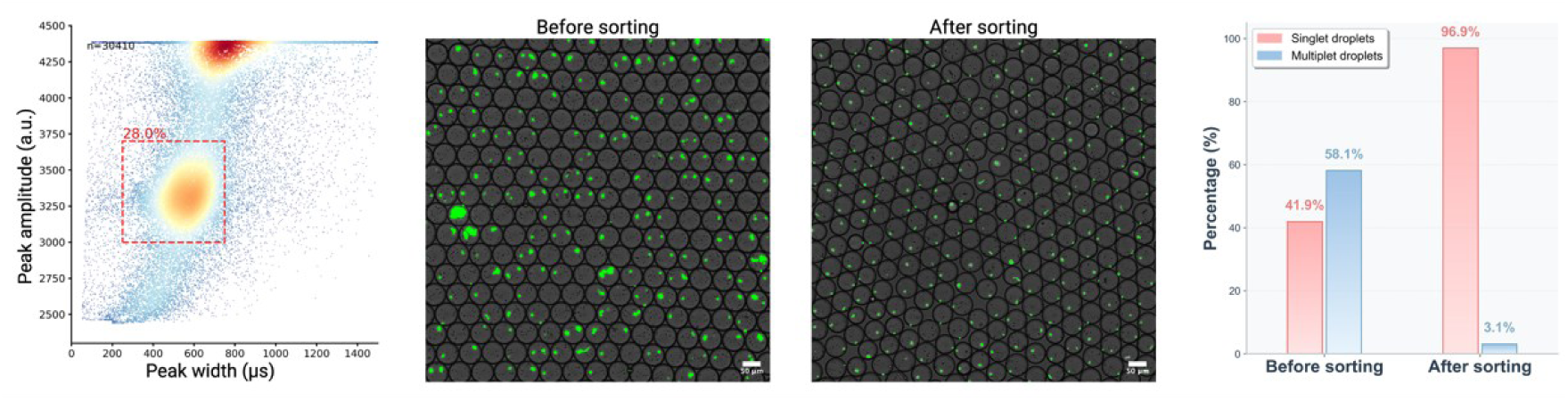
Sorting of Singlet Droplets for High Input Nuclei Concentration. **Left**: The amplitude-width scatter plots from 30,410 peaks with Poisson parameter λ ∼ 0.95. **Middle**: The representative microscopic images of droplets encapsulating nuclei before and after sorting. **Right**: The percentage of singlet and multiplet droplet before and after sorting. The percentage of singlet droplets significantly increased from 41.9% to 96.9% after sorting, while multiplet droplets decreased from 58.1% to 3.1%.

**Fig. S10.**
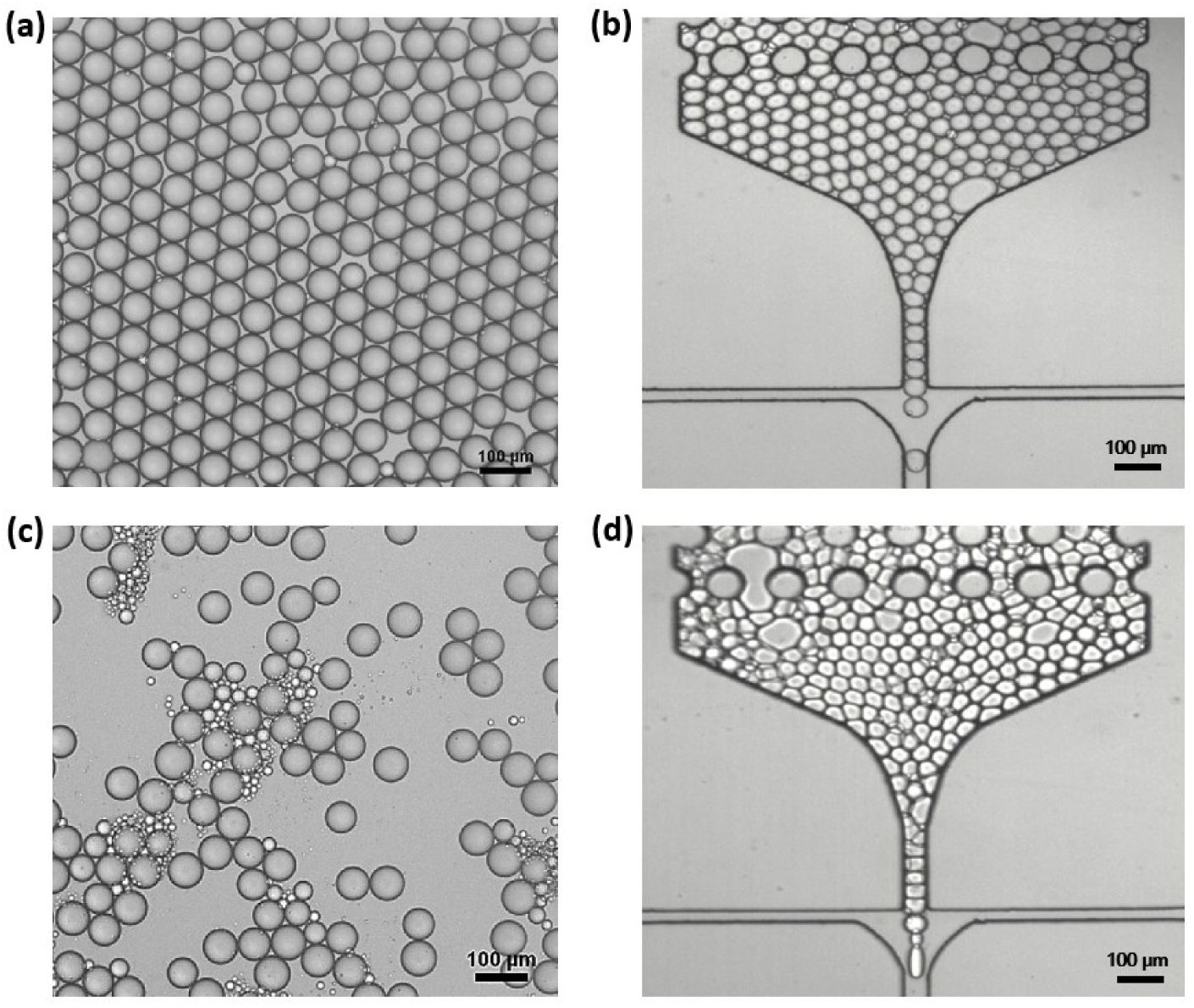
Different Reagents (proteinase K and SDS) Effect on Stability of Nuclei Droplets. (a, b) Nuclei droplets encapsulating proteinase K under 37℃ for 20 minutes and 55℃ for 15 minutes incubation before and after injection into the triple-droplet merging chip. (c, d) Nuclei droplets encapsulate SDS under 37℃ for 20 minutes incubation before and after injection into the triple-droplet merging chip.

**Fig. S11.**
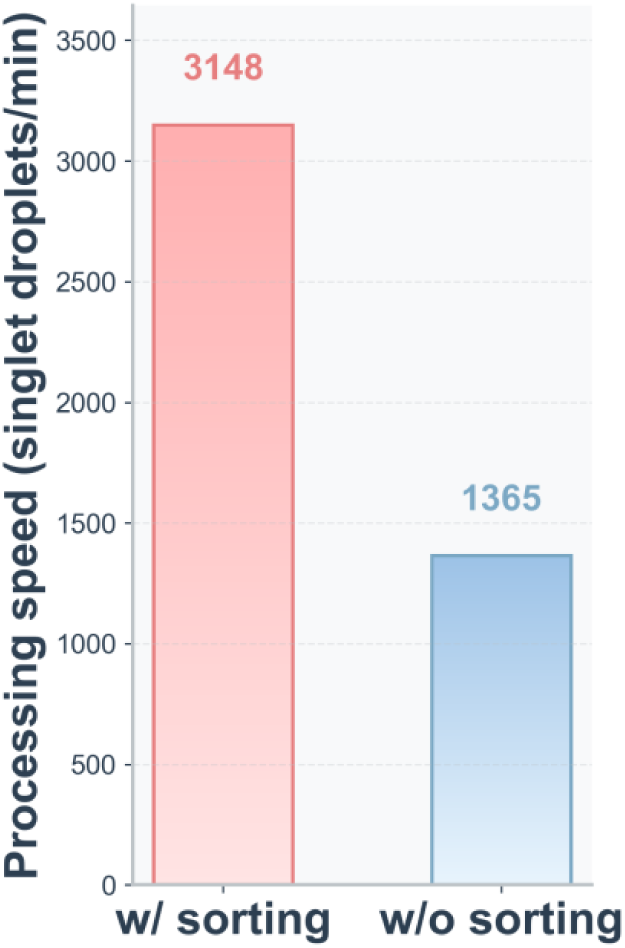
Processing Speed of Singlet Droplets with Sorting and without Sorting. The processing speed was calculated by the singlet droplet percentage, the barcode droplet positive rate and the triple-droplet triad rate comparing those with multi-parametric sorting and without.

**Fig. S12.**
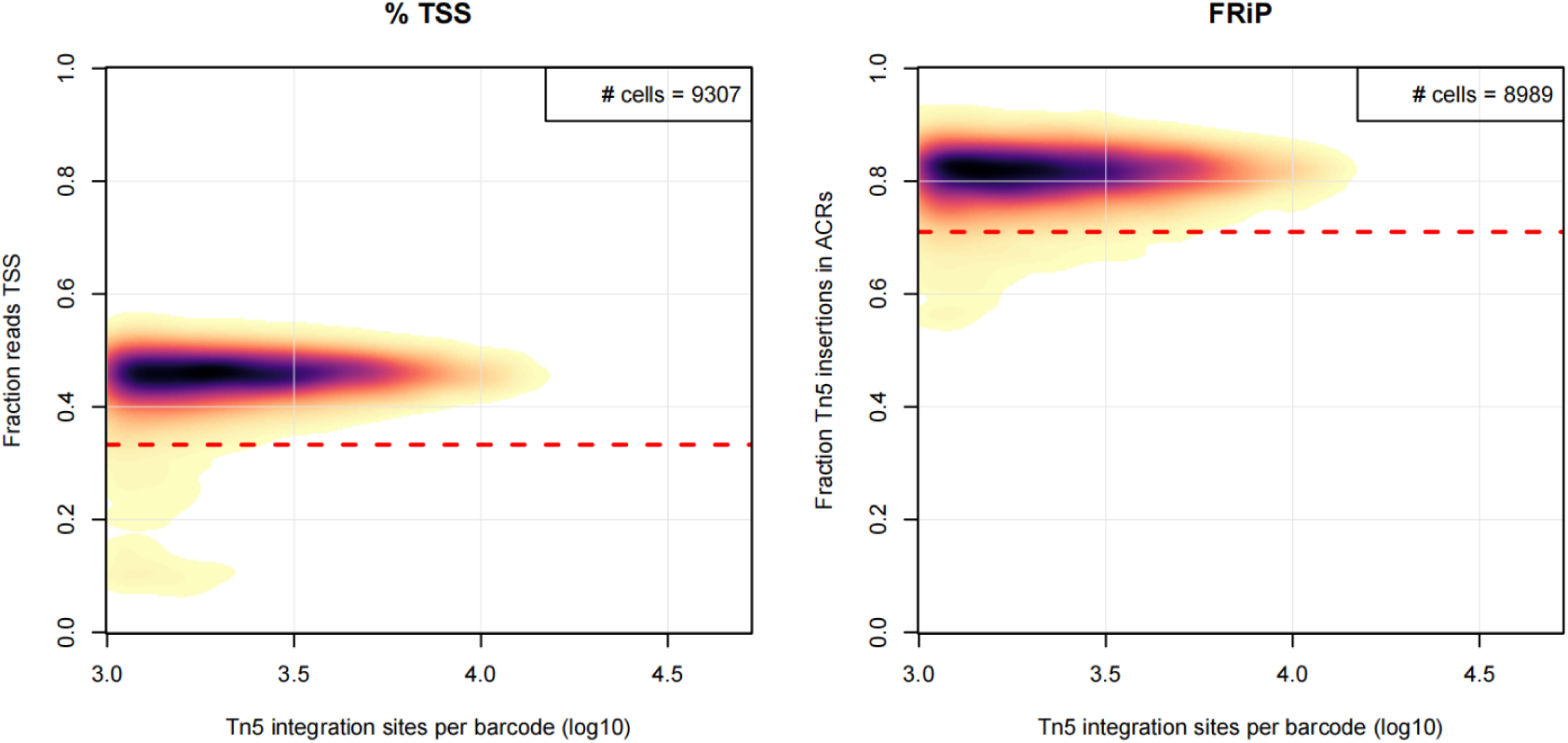
Quality Control Analysis of sc-MusTer-ATAC-seq Data using R Package Socrate. (a) Density plot showing the proportion of Tn5 sites located near the transcription start site (TSS) relative to the total reads for each cell barcode. (b) Density plot illustrates the number of Tn5 sites found in accessible chromatin peaks compared to the total number of Tn5 sites per cell barcode. The peaks were identified using the pseudo-bulked library.

**Fig. S13.**
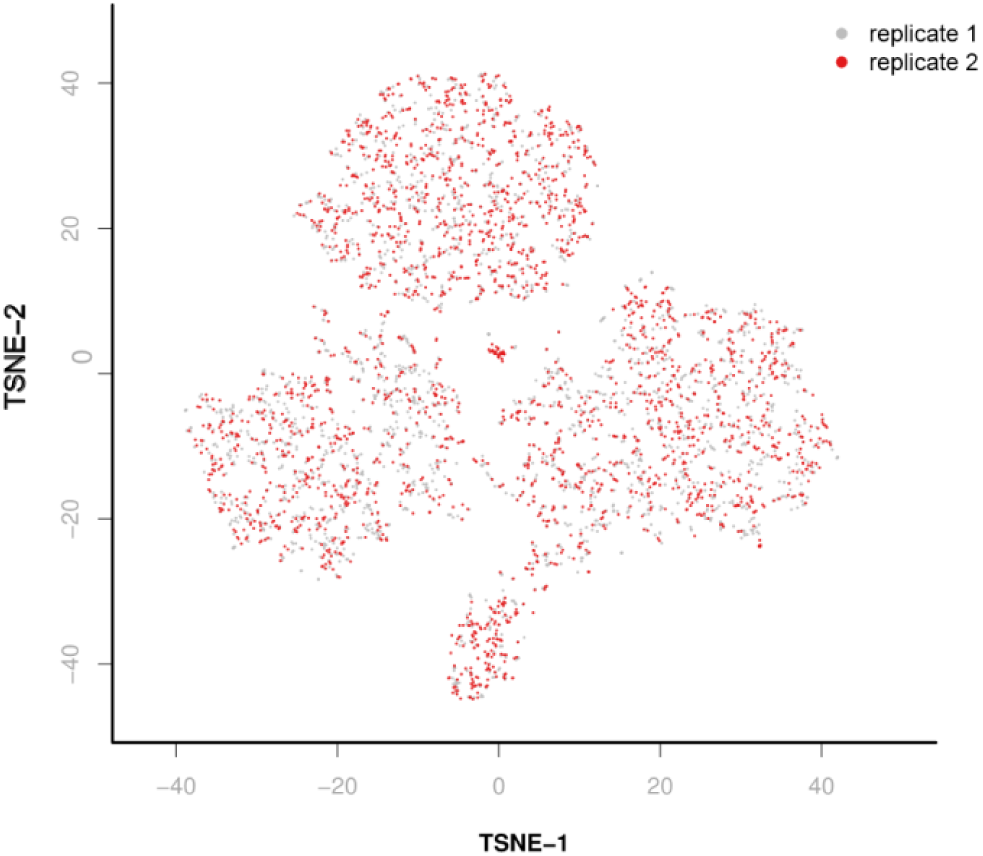
Two sc-MusTer-ATAC-seq Replicates for Clustering. The clustering of the two samples was highly similar, suggesting good reproducibility of our scATAC-seq method with MusTer.

**Fig. S14.**
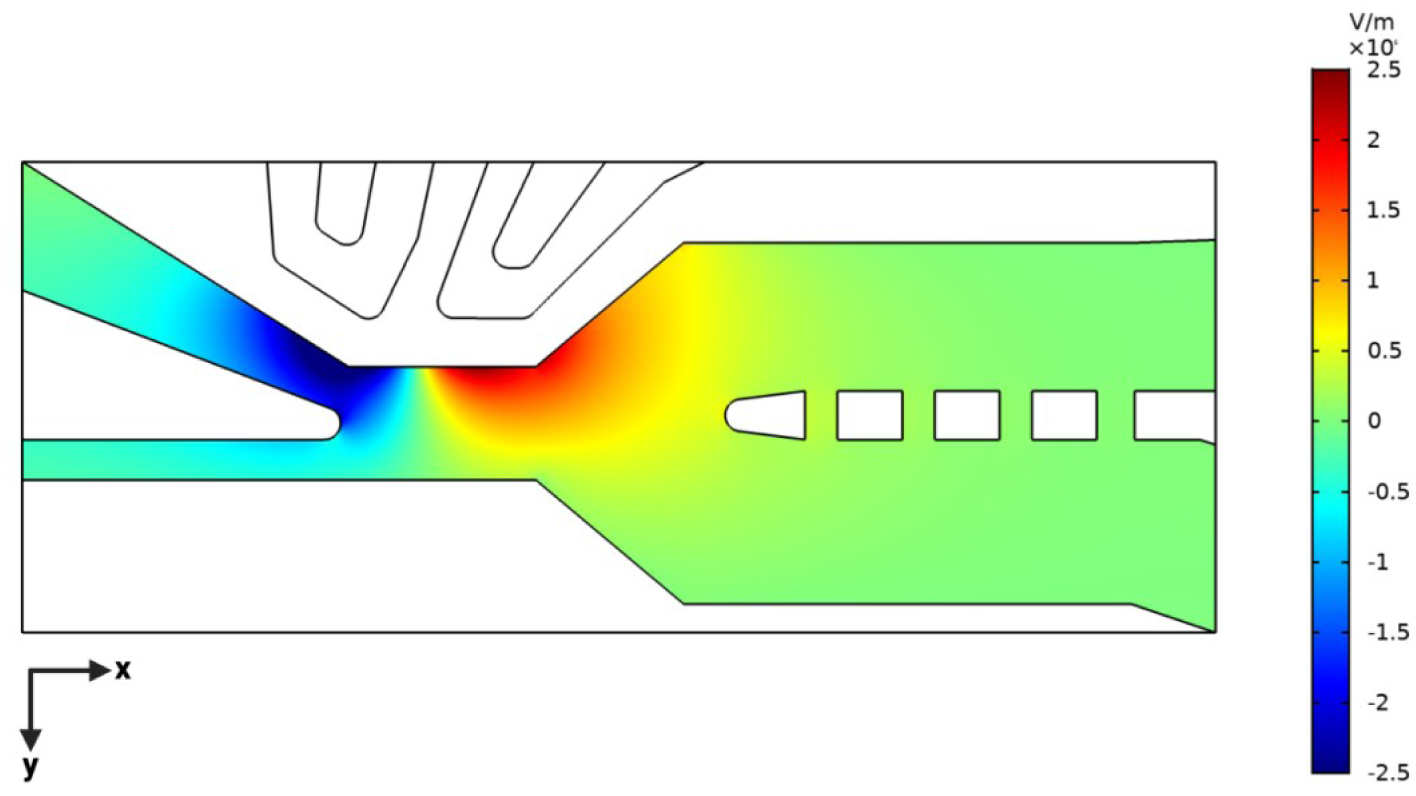
Simulated Distribution of Electric Field Intensity Along the y-axis at the Sorting region. The simulation was conducted using the Electrostatics interface of the AC/DC module in COMSOL Multiphysics^®^ v4.3 to model the electric field generated during droplet sorting. An alternating current (AC) voltage of 800 V at 20 kHz was applied to the sorting electrodes. All domains were set to satisfy charge conservation conditions. Material properties were defined using the relative permittivity of Novec™ 7500 for the channel (fluid) domain and PDMS for the surrounding channel walls. The resulting field distribution illustrated the spatial profile of electric field strength in the vertical (y) direction across the channel, governing the dielectrophoretic force applied to droplets during sorting.

**Fig. S15.**
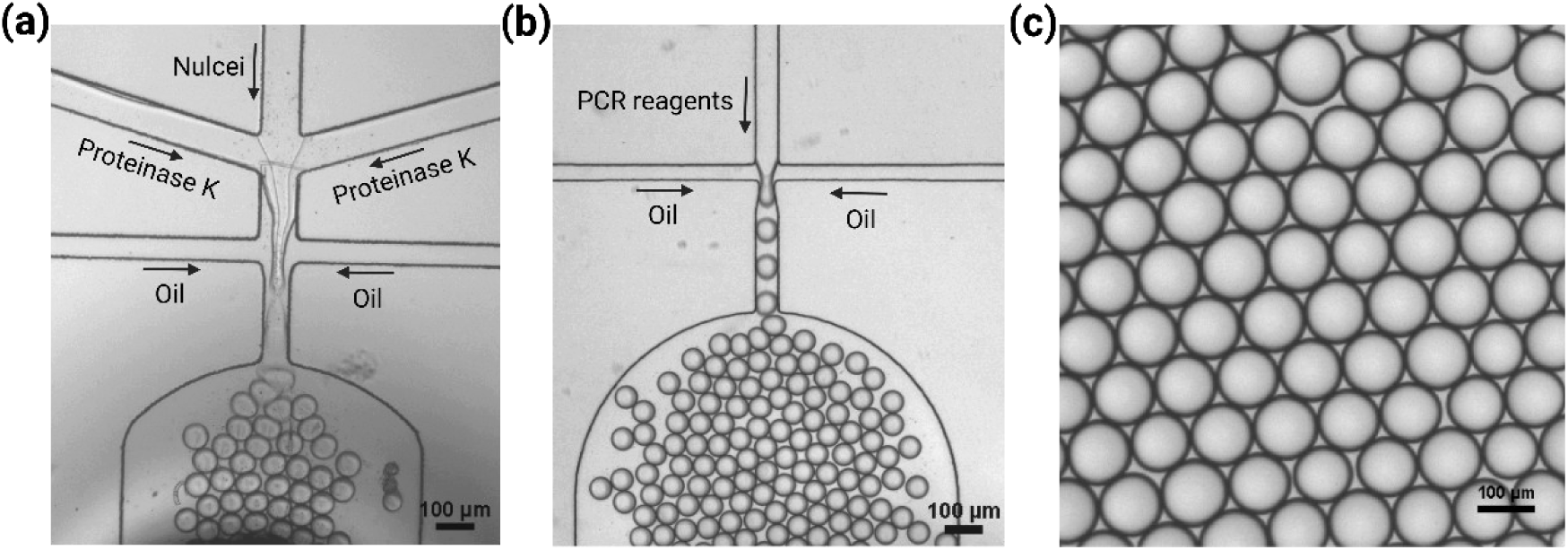
Nuclei Droplets, Barcode Droplets Generation and Droplets after Triple-droplet Merging. (a) A representative microscopy image of nuclei droplets generation process with the nuclei droplet generator. The size of generated nuclei droplets was about 64 µm in diameter. (b) A representative microscopy image of barcode droplets generation by the barcode droplet generator. The size of generated barcode droplets was about 50 µm in diameter. (b) A representative microscopy image of merged droplets after triple-droplet merging with the triple-droplet merging chip. The size of merged droplets was about 112 µm in diameter.

**Fig. S16.**
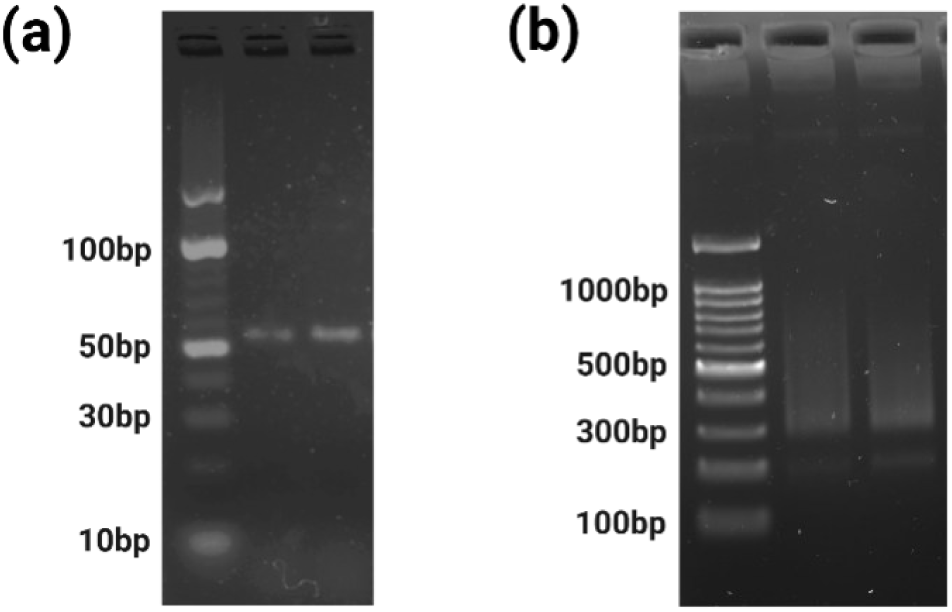
Gel Electrophoresis for Barcode Products and Sequencing Library. (a) Agarose gel electrophoresis (5%) of barcode products (53 bp) after 30 cycles ddPCR using 10 bp DNA ladder. (b) Agarose gel electrophoresis (1.5%) of final library for sequencing (300 - 800 bp) using 100 bp DNA ladder.

**Supplementary Movie 1.**

Characterization of singlet droplets and the four types of multiplet droplets.

**Supplementary Movie 2.**

Exclusion of Type IV multiplet-mimicking droplets encapsulating 10-µm fluorescent PS beads.

**Supplementary Movie 3.**

Exclusion of Type I-III multiplet-mimicking droplets encapsulating multiple 4-µm PS beads.

**Supplementary Movie 4.**

Exclusion all types of multiplet-mimicking droplets encapsulating 10-µm PS beads and multiple 4-µm PS beads.

**Supplementary Movie 5.**

Exclusion multiplet droplets encapsulating multiple tagmented nuclei.

**Supplementary Movie 6.**

On-chip PCR droplets generation and pairing with reinjected singlet droplets.

**Supplementary Movie 7.**

Barcode droplets, sorted nuclei singlet droplets and PCR droplets pairing.

**Supplementary Movie 8.**

Barcode droplets, sorted nuclei singlet droplets and PCR droplets merging.

**Supplementary Movie 9.**

Integration of nuclei droplets generation and sorting on a single microfluidic chip.

## Notes

### Competing Interest Statement

The authors have declared no competing interest.

